# Metabolic balancing by miR-276 shapes the mosquito reproductive cycle and *Plasmodium falciparum* development

**DOI:** 10.1101/548784

**Authors:** Lena Lampe, Marius Jentzsch, Elena A Levashina

## Abstract

*Anopheles* mosquitoes are obligate vectors of the human malaria parasite *Plasmodium falciparum*. The blood-feeding behavior of *Anopheles* females delivers essential nutrients for egg development and drives transmission from one human host to another. *Plasmodium* growth is adapted to the vector reproductive cycle, but how changes in the reproductive cycle impact parasite development is poorly understood. Here, we show that the blood meal-induced miR-276-5p fine-tunes the duration of the mosquito reproductive cycle. Silencing of miR-276 prolonged amino acid catabolism and increased female fertility rates, suggesting that timely termination of the reproductive cycle restricts mosquito investment into reproduction. Prolongation of the reproductive period in *P. falciparum*-infected females compromised the development of the transmissible parasite form called sporozoite. Our results suggest that *Plasmodium* sporogony exploits surplus resources after mosquito reproductive investment and demonstrate the crucial role of the mosquito amino acid metabolism in parasite within-vector proliferation and malaria transmission.

## Introduction

The hematophagous lifestyle of *Anopheles* females is an efficient reproductive strategy that is exploited by the malaria parasite for its transmission. Mosquitoes mostly feed on carbohydrate-rich plant nectars and only females take blood to boost reproduction. The blood meal provides the female with a nutritional boost in amino acids and lipids that is crucial for egg development. This extreme change in diet triggers massive coordinated metabolic changes in multiple mosquito tissues to ensure robust egg development within a three-day reproductive cycle.

The insect fat body, the main nutrient storage organ, plays a central role in the mosquito metabolism. During a pre-blood-feeding anabolic phase, the fat body accumulates nutrients in the form of triacylglycerides (TAGs) and glycogen derived from plant nectars (Briegel et al., 2002; Hou et al., 2015; Wang et al., 2017; Ziegler and Ibrahim, 2001). A successful blood feeding massively increases levels of free amino acids (AAs) in the mosquito circulation, that are sensed in the fat body by the target of rapamycin (TOR) pathway (Carpenter et al., 2012; Hansen et al., 2005; Roy et al., 2015). Activation of TOR, together with the increasing levels of the steroid hormone 20-hydroxyecdysone (20E), initiate a switch from anabolic to catabolic metabolism (Hou et al., 2015; Roy et al., 2015; Wang et al., 2017). In the fat body, blood meal-derived AAs feed into the tricarboxylic acid (TCA) cycle to generate energy for nutrient mobilization and are incorporated into major yolk proteins such as lipophorin and vitellogenin (Zhou et al., 2004a, 2004b). Inhibition of nutrient transport, metabolism, or the TOR signaling pathway dramatically reduces egg production (Carpenter et al., 2012; Fuchs et al., 2014a; Gulia-Nuss et al., 2011; Hansen et al., 2004; Rono et al., 2010). Therefore, the mosquito metabolic program during the early (0-10 h post blood feeding (hpb)) and mid-phase (10-36 hpb) of the reproductive cycle prioritizes the mobilization of nutrients for rapid egg development. During the late phase (36–72 hpb), production of the yolk proteins in the fat body ceases, and transcriptional and metabolic programs return to the anabolic state to replenish consumed glycogen and lipid reserves (Hou et al., 2015; Wang et al., 2017). The metabolic switch from catabolism to anabolism during the late phase prepares females for the next reproductive cycle. Indeed, prolongation of the reproductive cycle by inhibition of autophagy or silencing of a negative regulator of TOR signaling in the fat body, curbs egg production in the second reproductive cycle (Bryant et al., 2011; Mane-Padros et al., 2012a). Therefore, both the induction and termination of the reproductive cycle are essential for female fertility.

Mosquitoes acquire malaria parasites when feeding on blood infected with *P. falciparum* sexual stages of that fuse in the mosquito midgut to produce motile ookinetes. The ookinetes traverse the midgut epithelium at 18-24 h post infectious blood feeding (hpb) and transform into oocysts to establish infection at the basal side of the midgut wall. Here, the parasites undergo massive replication and maturation (also called sporogony). Within two weeks of sporogony, the parasite biomass increases dramatically generating up to 1,000 sporozoites within each oocyst (Stone et al., 2013). Developing oocysts require large amounts of nutrients such as AAs, sugars and lipids, for which they rely on the mosquito environment. Therefore, it is not surprising that the mosquito’s nutritional status largely correlates with parasite development (Moller-Jacobs et al., 2014; Takken et al., 2013). Inhibition of sugar uptake or lipid access to the *Plasmodium* oocysts decreases the number and virulence of transmissible sporozoites (Costa et al., 2018; Slavic et al., 2011). As sporogony occurs after completion of the mosquito reproductive cycle, the oocysts do not directly compete with the vector but rely on the unconsumed blood meal-derived lipids stored in the mosquito tissues (Costa et al., 2018).

Hormonal and amino acid signals tightly control transcriptional and post-transcriptional network that shapes mosquito metabolism across multiple tissues. In this network, microRNAs contribute to post-transcriptional tuning and link the endocrine regulation with metabolic homeostasis in *Aedes* mosquitoes (Liu et al., 2014; Lucas et al., 2015). However, how the miRNAs that regulate mosquito metabolism impact vector-parasite interactions is unknown. In *A. gambiae*, a major malaria vector in sub-Saharan Africa, blood feeding increases the fat body levels of miR-276-5p transcripts (miR-276 hereafter) during the mid-/late-phase of the reproductive cycle (Lampe and Levashina, 2018). Here, we report that miR-276 tunes the duration of the reproductive cycle by inhibiting expression of the *branched chain amino acid transferase* in the fat body. We show that miR-276 depletion prolongs amino acid catabolism, thereby benefiting female fertility. Prolongation of the reproductive cycle compromises *P. falciparum* sporogonic development and reduces the number of transmissible sporozoites. Our results demonstrate the important role of mosquito amino acid metabolism in vector competence and malaria transmission.

## Results

### Amino acid and steroid hormone signaling regulate blood feeding-induced expression of miR-276 in the mosquito fat body

To identify regulators of the mosquito reproductive cycle, we examined the expressional profile of the mature miR-276 during the first reproductive cycle by reverse transcription quantitative real-time PCR (RT-qPCR) in the fat body samples collected before and after blood feeding. Newly-eclosed females showed high levels of miR-276 expression that declined on the second day after eclosion (Fig. S1). A blood meal transiently increased miR-276 levels from 28 to 44 hpb (Fig. 1A). This expression pattern paralleled the kinetics of the steroid hormone ecdysone, a key regulator of mosquito development and reproduction, whose synthesis is triggered by a blood meal. To compare the timing of miR-276 expression and ecdysone titers, we measured the levels of 20-hydroxyecdysone (20E), the metabolically active form of ecdysone, in the blood-fed females using ELISA (Fig. 1A). 20E titers preceded the induction of miR-276 expression as they increased from 6 to 24 hpb and fell down to basal levels at 36 hpb. Based on these results, we hypothesized that expression of miR-276 in the fat body may be regulated by 20E.

**Figure 1:**
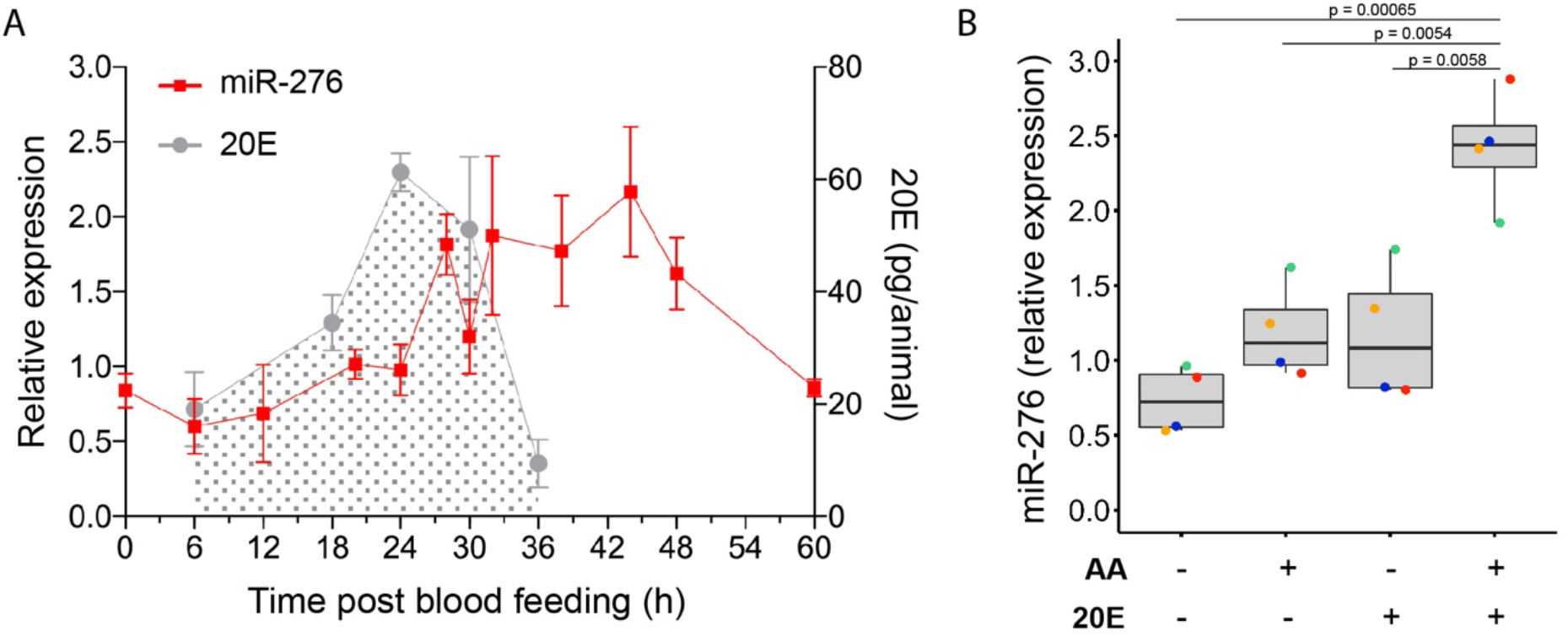
Amino acid and steroid hormone signaling regulate expression of miR-276 in the fat body of blood-fed mosquitoes. (A) miR-276 (red) expression in the female fat body (n=10, N=6) and 20-hydroxyecdysone (20E) titres (grey) in the mosquito females (n=9, N=3) at different time points after blood feeding. The plots show mean ± SEM of independent experiments. (B) miR-276 expression in *ex vivo* fat body cultures of 3-4-day-old females (n=3, N=4) after incubation with medium with or without amino acids (AA) and 20E. miRNA expression levels were normalized using the ribosomal protein *RPS7* gene. Boxplots show the median with first and third quartile, whiskers depict the min and max of independent experiments, coloured dots show mean of each experiment. The statistical significance was tested by one-way ANOVA followed by Tukey’s post hoc test and the obtained p-values are shown above the horizontal lines. n = number of mosquitoes pooled for each independent experiment; N = number of independent experiments.

To identify the triggers of miR-276, we gauged its expression levels by RT-qPCR in an *ex-vivo* fat body culture system after stimulation with 20E or amino acids (AA) (Chung et al., 2017; Deitsch et al., 1995). The fat body tissues were incubated in culture medium supplemented or not with 20E and/or AA *in vitro*. Addition of AA or 20E increased miR-276 levels by two-fold. Moreover, a significant five-fold induction of miR-276 expression was observed when AA and 20E were added together (Fig. 1B). These results suggested a synergistic effect of AA and 20E on regulation of miR-276 expression in the fat body after a blood meal.

### miR-276 inhibits expression of the branched chain amino acid transferase

MicroRNAs post-transcriptionally promote transcript degradation or inhibition of translation by binding to specific sites located predominantly in the 3’UTR. To identify putative miR-276 mRNA targets, we applied three *in silico* target prediction tools, namely MiRanda (John et al., 2004), RNAhybrid (Kruger and Rehmsmeier, 2006) and MicroTar (Thadani and Tammi, 2006)(Table S1). We next used a publicly available RNAseq dataset (Roy et al., 2015) to identify the fat body-expressed candidate target mRNAs with the reversed to miR-276 expression pattern. This dual approach allowed us to down-select eight putative targets for further analyses: *ATP synthase* (AGAP012081), *branched chain amino acid transferase* (*BCAT*; AGAP000011), *glycerol-3-phosphate dehydrogenase* (*GPD*; AGAP004437), *methylmalonate-semialdehyde dehydrogenase* (*MMSDH*; AGAP002499), *NADH dehydrogenase* (*NADH (I)*, AGAP010464), *3-methylcrotonyl-CoA carboxylase* (*MCC*; AGAP010228), *branched-chain alpha-keto acid dehydrogenase* (*BCKDH*, AGAP003136) and *mitochondrial ornithine receptor* (*MOR;* AGAP000448).

We posited that expression of the target transcripts should be repressed by miR-276 and measured transcript levels of the selected mRNAs in the fat bodies of miR-276 knockdown mosquitoes (miR-276^KD^) at the peak of miR-276 expression (38 hpb) by RT-qPCR. miR-276 was silenced by injection of antisense oligonucleotides (miR-276 antagomirs), whereas injection of a scrambled antagomir served as control (Table S3). Out of the examined candidates, only expression of *BCAT* significantly increased after miR-276 inhibition (Fig. 2A). *BCAT* carries one predicted miR-276 binding site in its 3’UTR (Fig. 2B). However, as we did not perform genome-wide analysis of mRNA changes following miR-276 inhibition, we cannot exclude that miR-276 regulates other mRNAs than *BCAT*.

**Figure 2:**
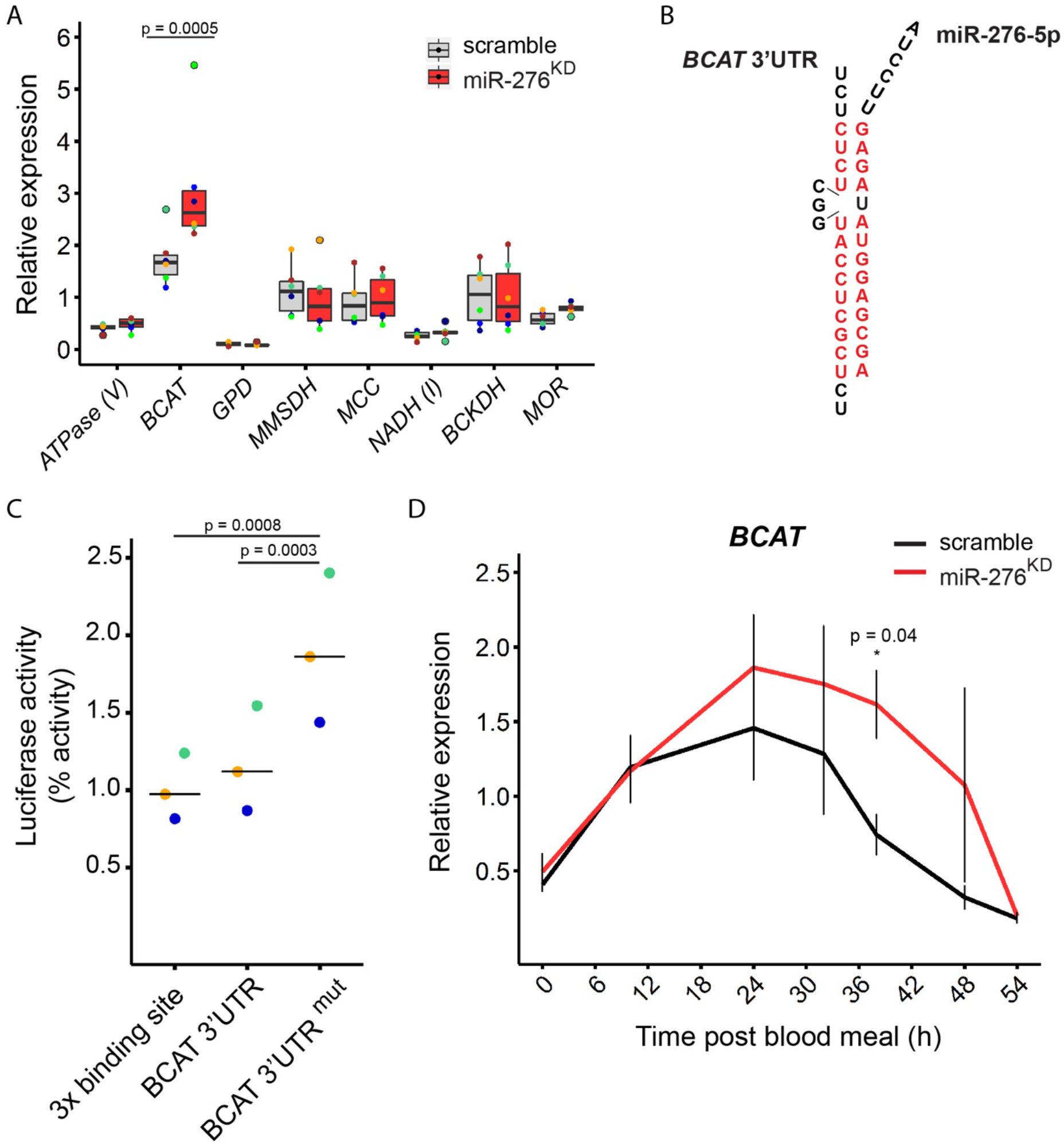
miR-276 post-transcriptionally represses the *branched chain amino acid transferase* in the mosquito fat body. (A) Expression of the predicted miRNA targets at 38 h post blood feeding in the fat body of mosquitoes (n=5, N=6) injected with anti-miR-276 (miR-276, red) or scrambled antagomir (scramble, grey). Expression of the *ATP synthase (ATPase V), branched chain amino acid transferase* (*BCAT*), *Glycerol-3-phosphate dehydrogenase* (*GPD*), *methylmalonate-semialdehyde dehydrogenase* (*MMSDH*), *3-methylcrotonyl-CoA carboxylase* (*MCC*), *NADH dehydrogenase* (*NADH (I)*), *branched-chain alpha-keto acid dehydrogenase* (*BCKDH*) and *mitochondrial ornithine receptor* (*MOR*). Expression levels were normalized using the ribosomal protein *RPS7* gene. Boxplots show the median with first and third quartile, whiskers depict the min and max. Statistical significance was tested by two-way ANOVA followed by Tukey’s *post hoc* test and significant differences are shown by the *p*-value above the horizontal line. (B) Sequence of the predicted miR-276 binding site in the *BCAT* 3’UTR. Red color indicates complementary nucleotides in the 3’UTR binding site and miR-276. (C) Dual luciferase reporter assay *in vitro* showing that miR-276 directly targets the 3’-UTR of *BCAT*. A construct containing three copies of miR-276 binding sites served as positive control. Lines show the median and colored dots show means of independent experiments (N=3). Statistical significance of differences was tested by one-way ANOVA followed by Tukey’s post hoc test and significant differences are indicated by the *p*-values above the horizontal lines. (D) *BCAT* expression in the fat body during the reproductive cycle of mosquitoes injected with anti-miR-276 (miR-276^KD^, red) or scrambled antagomir (scramble, black). Expression levels were normalized using the ribosomal protein *RPS7* gene. Means ± SEM (vertical lines) were plotted. Statistically significant differences between miR-276^KD^ and control mosquitoes were determined by two-way ANOVA followed by Tukey’s *post hoc* test (N=3) and significant differences are shown by the *p*-value (*) (n = number of mosquitoes pooled for each independent experiment; N = number of independent experiments).

We next tested for direct interaction between miR-276 and the binding site in the *BCAT* 3’UTR using a Dual Luciferase Reporter Assay in the *Drosophila* Schneider 2 (S2) cell cultures *in vitro*. This assay examines the ability of miR-276 to post-transcriptionally inhibit a reporter construct containing the *Renilla* luciferase reporter gene fused to the *BCAT* 3’UTR. We generated three constructs: (1) a positive control, which carries three copies of the miR-276 binding site (3x binding site), (2) the wild-type *BCAT* 3’UTR with an intact single miR-276 binding site (BCAT 3’UTR); and (3) a *BCAT* 3’UTR negative control with the mutated sequence of miR-276 binding site (BCAT 3’UTR^mut^). S2 cells expressed low levels of endogenous *Drosophila* miR-276 leading to basal inhibition of the *BCAT* 3’UTR reporter activity. Co-transfection of the mosquito miR-276 further enhanced inhibition of the luciferase reporter fused to the functional 3’UTR binding sites but not to the mutated control (Fig. S2). Importantly, mosquito miR-276 significantly repressed expression of the positive control with the 3 copies of the miRNA binding site and the wild-type *BCAT* 3’UTR. In contrast, mutation of the BCAT miRNA binding site abolished the inhibition of the luciferase reporter by more than two-fold (Fig. 2C and Fig. S2). These results demonstrated that *BCAT* 3’UTR is a *bona fide* target of miR-276.

To confirm our *in vitro* findings, we examined *BCAT* transcript levels *in vivo* after blood feeding in miR-276^KD^ and scramble-injected mosquitoes. We observed that in the presence of miR-276, *BCAT* transcript levels went down as early as 24 hpb, inhibition of the miRNA function by antagomir prolonged *BCAT* expression by almost 12 h (Fig. 2D). We concluded that miR-276 tunes down *BCAT* expression during the late phase of the mosquito reproductive cycle.

### miR-276 fine tunes amino acid metabolism during the late phase of the reproductive cycle

BCAT catalyzes the first step of the branched chain amino acid (Leucine, Valine and Isoleucine) catabolism. As miR-276 post-transcriptionally repressed *BCAT* during the late phase of the reproductive cycle, we examined the amino acid metabolism after miR-276 knockdown. We compared metabolite profiles of miR-276^KD^ and control females (whole mosquitoes) at three time points after *P. falciparum*-infected blood feeding (10, 38 and 48 hpb) using gas chromatography - mass spectrometry (GC-MS) and liquid chromatography - mass spectrometry (LC-MS). We selected time points before (10 hpb) and during miR-276 expression (38 and 48 hpb). In total, 4,716 chromatographic peaks (*m/z* at a specific retention time; 816 annotated) were detected, of which 2,426 features were included in the heatmap after IQR-based filtering (Fig.3A). Massive changes in metabolites during the reproductive cycle were detected in both control and miR-276^KD^ mosquitoes. At the early as 10 hpb, a large cluster of highly enriched metabolites correlated with the influx of human blood. This cluster decreased with blood digestion (38 hpb) and disappeared at the late phase of the reproductive cycle (48 hpb). Conversely, other metabolite clusters increased with blood digestion and ovary development (Fig. 3A). Overall, the observed metabolite dynamics was not significantly perturbed by miR-276^KD^ and closely recapitulated the major physiological processes induced by blood feeding. This observation was in line with the fine-tuning role of miR-276, whose silencing did not hinder major blood feeding-induced metabolic changes.

**Figure 3:**
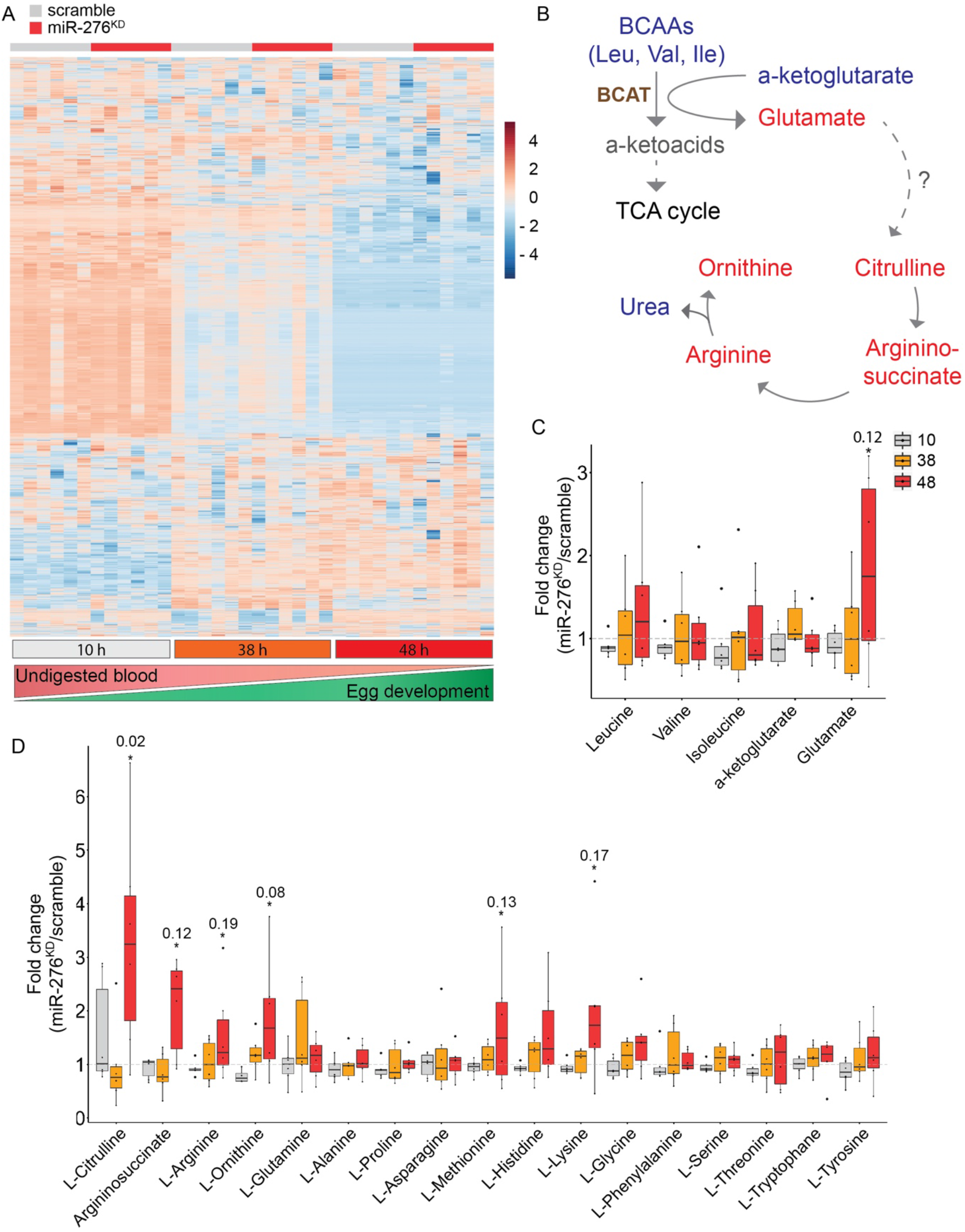
miR-276 fine-tunes AA metabolism in the late phase of the mosquito reproductive cycle. (A) Heatmap of metabolite levels measured by GC-MS and LC-MS from whole females (n=10, N=6) at 10, 38 and 48 h post *P. falciparum*-infected blood meal in control (scrambled, grey) and miR-276-depleted (miR-276^KD^, red) mosquitoes. Bottom bars indicate time points after a blood feeding, and the gradients below show the progression of blood digestion and ovary development. Metabolites are plotted on the y-axis and samples on the x-axis. For heatmap visualization, data was log2 transformed, mean-centered and divided by the square root of each metabolite. (B) Overview of the classical enzymatic activity of the branched chain aminotransferase (BCAT) in animals. The amino group of BCAAs is transferred to α-ketoglutarate by BCAT forming glutamate. Glutamate serves as a nitrogen sink, which is normally passed on to the urea cycle in vertebrates. As mosquitoes lack the carbamoyl phosphate synthase, the link between glutamate and citrulline of the urea cycle is unknown. Within the classical urea cycle, citrulline is converted to argininosuccinate and subsequently arginine, which is finally metabolized to ornithine releasing urea. Metabolites changed upon miR-276 inhibition (48 hpb) are in red, metabolites at constant levels are in blue, and undetected metabolites are in grey. (C) Fold-change differences in branched chain amino acids (leucine, valine, isoleucine), α-ketoglutarate and glutamate detected in miR-276^KD^ as compared to scramble controls. (D) Fold change differences in metabolites of AA metabolism at 10 h (grey), 38 h (orange), 48 h (red) post blood feeding in whole females. Statistical significance of differences in metabolite levels of miR-276^KD^ and scramble controls was tested by t-test, n = number of mosquitoes pooled for each independent experiment; N = number of independent experiments.

We next focused on BCAT-related metabolites. BCAT initiates BCAA catabolism by transferring the BCAA amino group to α-ketoglutarate forming glutamate, which is thought to serve as a nitrogen sink in insects (Scaraffia et al., 2006) (Fig. 3B). While BCAA levels did not massively change in miR-276^KD^ mosquitoes, significantly higher levels of glutamate were detected at 48 hpb (Fig. 3C). Furthermore, miR-276^KD^ also increased levels of histidine, lysine, citrulline, arginine, argininosuccinate and ornithine at 48 hpb (Fig. 3D), suggesting an accumulation of metabolites associated with nitrogen metabolism, the waste product of AA catabolism, during the late phase of reproductive cycle. The detected increase in nitrogen metabolism in miR-276^KD^ mosquitoes specifically during the late phase of the reproductive cycle (48 hpb) is consistent with the transcriptional increase in *BCAT* levels observed in miR-276^KD^ females. Taking together, our results suggest that miR-276 contributes to the termination of the AA catabolism by fine-tuning *BCAT* transcript levels and that miR-276 silencing prolongs the catabolic phase of the mosquito reproductive cycle.

### Prolongation of the reproductive cycle by miR-276 silencing unleashes mosquito metabolic investment into oogenesis and compromises P. falciparum sporogonic development

Both mosquitoes and malaria parasites rely on nutrients for successful ovary and sporogonic development, respectively. We first examined how prolongation of the catabolic phase in miR-276^KD^ mosquitoes affects oogenesis. Female mosquitoes were injected with the miR-276 or control (scramble) as described above and blood fed three days later to induce egg development. Two days later, females were transferred to individual egg laying chambers and the numbers of eggs laid by individual females were enumerated. Inhibition of miR-276 significantly increased egg laying (fertility) without compromising larval hatching rates (fecundity) (Fig. 4A and Fig. S3). We concluded that miR-276 function restrains mosquito metabolic investment into oogenesis by terminating the catabolic phase before complete nutrient consumption by oogenesis.

**Figure 4:**
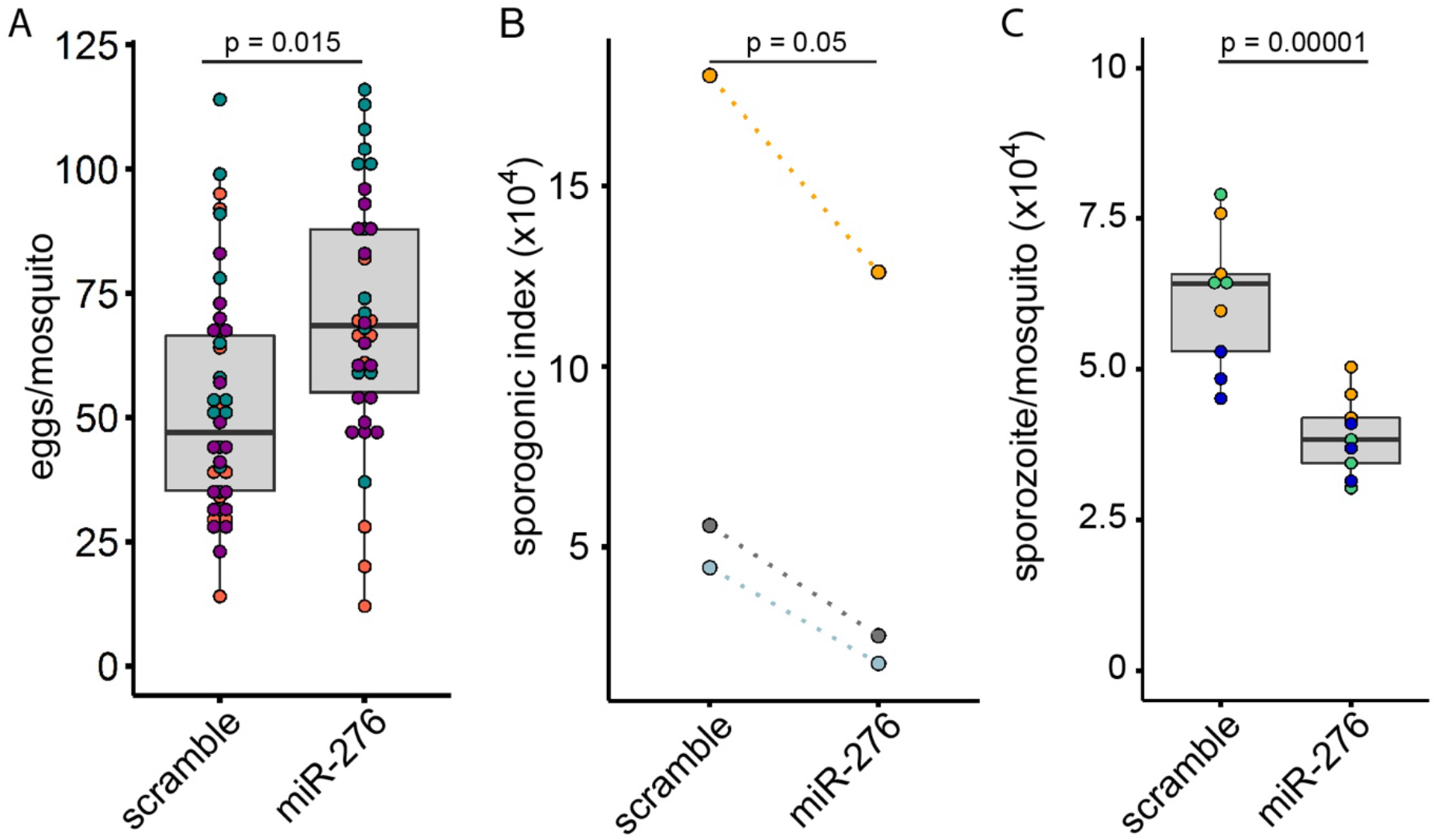
Effects of miR-276 silencing on mosquito fertility and *P. falciparum* sporogonic development. (A) Egg laying rates (fertility) of individual females injected with miR-276 (miR-276^KD^) or control antagomir (scramble) after a blood meal. Boxplots show the median with first and third quartile, whiskers depict min and max values. Each dot represents one mosquito and dot colors indicate independent experiments (N=3, Table S4). (B) Females injected with miR-276 (miR-276^KD^) or control (scramble) antagomir were infected with *P. falciparum*. At day 11 post infection, oocyst number and size were measured. The median of *P. falciparum* oocysts per midgut and mean of *P. falciparum* oocyst size per experiment were multiplied to generate a sporogonic index (N=3, Table S4). (C) The numbers of the salivary gland sporozoites per mosquito were quantified on day 14 post infection. Boxplots show the median with first and third quartile, whiskers depict min and max values. Colors show replicates of each independent experiment. Statistically significant differences are indicated by *p*-values above the horizontal lines deduced by one-way ANOVA and Tukey’s post-hoc test (N=3, Table S4), N = number of independent experiments.

We asked whether the observed shift in the metabolic investment into oogenesis impacts parasite development. Using the same experimental settings, we infected miR-276^KD^ and control mosquitoes with *P. falciparum* and gauged the number and size of oocysts 11 days post infection. Inhibition of miR-276 significantly decreased either the median number or the mean size of *P. falciparum* oocysts. To quantify oocyst development, we calculated a sporogonic index by multiplying the median number by the mean size of oocysts per midgut in each independent experiment. We observed that regardless of infection levels, the sporogonic index was significantly lower in miR-276^KD^ mosquitoes than in controls (Fig. 4B). We further examined whether the observed decrease in the parasite mass translated into lower numbers of the salivary gland sporozoites. Indeed, miR-276 inhibition significantly reduced by two-fold the loads of the salivary gland sporozoites (Fig. 4C), suggesting that prolongation of the catabolic phase restricts resources available for the development of the parasite transmissible forms.

In summary, our results demonstrated that the miR-276-regulated switch from catabolic to anabolic amino acid metabolism in the fat body restricts mosquito investment into oogenesis and benefits *P. falciparum* development.

## Discussion

The hematophagous lifestyle equips mosquitoes with a speedy within-days egg development, but also benefits the transmission of the malaria parasite. Blood feeding induces massive metabolic changes that re-direct all resources towards reproduction, a state that is unsustainable over a long period of time. Here, we provide evidence that miR-276 contributes to a negative feedback loop that resets mosquito metabolism by repressing BCAA catabolism. This metabolic reset restricts mosquito reproductive investment and, thereby, benefits *P. falciparum* sporogony. Our results suggest that reproductive investment plays a crucial role in the mosquito vector competence.

The fat body is the major insect storage organ, which produces the majority of hemolymph vitellogenic proteins and critically contributes to mosquito ovary development (Arrese and Soulages, 2010). After blood feeding, high titers of AAs and 20E promote a metabolic switch in the fat body from anabolism to catabolism, resulting in a rapid release of glycogen and lipid reserves (Hou et al., 2015; Wang et al., 2017). The same signals induce expression of negative regulators, including miR-276, that warrant the timely reset of the fat body metabolism (Bryant et al., 2011; Lampe and Levashina, 2018; Mane-Padros et al., 2012b). Massive degradation of AAs in the fat body supports high energy demands of nutrient mobilization and protein synthesis for successful egg development (Fuchs et al., 2014b; Zhou et al., 2004a). However, high levels of AA degradation also generate a considerable amount of nitrogen waste, which may restrain the metabolic investment during the reproductive cycle (Isoe and Scaraffia, 2013; Isoe et al., 2017; Mazzalupo et al., 2016). Although *Aedes* mosquitoes lack the full repertoire of the urea cycle enzymes and rely on an alternative ammonia detoxification pathway engaging glutamine, alanine and proline (Scaraffia et al., 2006, 2010), we did not observe miR-276-driven changes in these metabolites at any tested time point. Instead, we detected increased levels of glutamate and other classic urea cycle metabolites (citrulline, ornithine, argininosuccinate and arginine) during the late phase of the reproductive cycle. These results point to potentially important metabolic differences between *Aedes* and *Anopheles* mosquitoes and call for further targeted metabolome investigations using N^15^-labelled BCAAs to identify the urea cycle components in both mosquito species. The timing and nature of metabolic changes observed in this study suggest that miR-276 contributes to fine regulation of AA catabolism in the late phase of the reproductive cycle. During this phase, the fat body undergoes a reverse shift from catabolic to anabolic metabolism in order to replenish nutrient resources in preparation for the next reproductive cycle (Hou et al., 2015; Wang et al., 2017). As part of this metabolic switch, miR-276 post-transcriptionally represses translation of the mRNA encoding the branched chain amino acid transferase, the catalyzer of the first step of BCAA breakdown. BCAA turnover represents a central hub that regulates the overall metabolic state and is tightly regulated (Li et al., 2017; She et al., 2007; Yoon, 2016; Zhang et al., 2017). In *Drosophila*, BCAA degradation is controlled by miR-277 (Esslinger et al., 2013). Interestingly, the BCAA degradation pathway in *Aedes* shapes the bacterial load in the midgut by an as yet unknown mechanism (Short et al., 2017). Therefore, in addition to the fat body examined here, post-transcriptional regulation of BCAA metabolism by miR-276 may affect mosquito physiology at multiple levels, including the midgut microbiome. Our results suggest that inhibition of miR-276 extends the reproductive cycle by prolonging BCAA catabolism and benefits female fertility. We propose that miR-276 contributes to a negative feedback loop that shifts the catabolic metabolism induced by AAs and 20E during the early-/mid-phase of the female reproductive cycle to the anabolic phase. Timely termination of the catabolic phase is important to restrict the costly reproductive investment and prevent complete nutrient exhaustion (Fig. 5A). Metabolic costs likely play an important role in the regulation of reproductive investment as blood digestion in hematophagous insects escalates oxidative stress and nitrogen waste products (Isoe et al., 2017; Magalhaes et al., 2008; Mazzalupo et al., 2016). Indeed, blood digestion reduces mitochondrial activity of the flight muscles, presumably to mitigate the overall metabolic stress (Gonçalves et al., 2009). Furthermore, timely conclusion of the reproductive program is essential in preparation for the next reproductive cycle (Bryant et al., 2011; Mane-Padros et al., 2012a). We propose that the duration of the reproductive cycle is tightly regulated by a balance between resource availability and reproductive costs.

**Figure 5:**
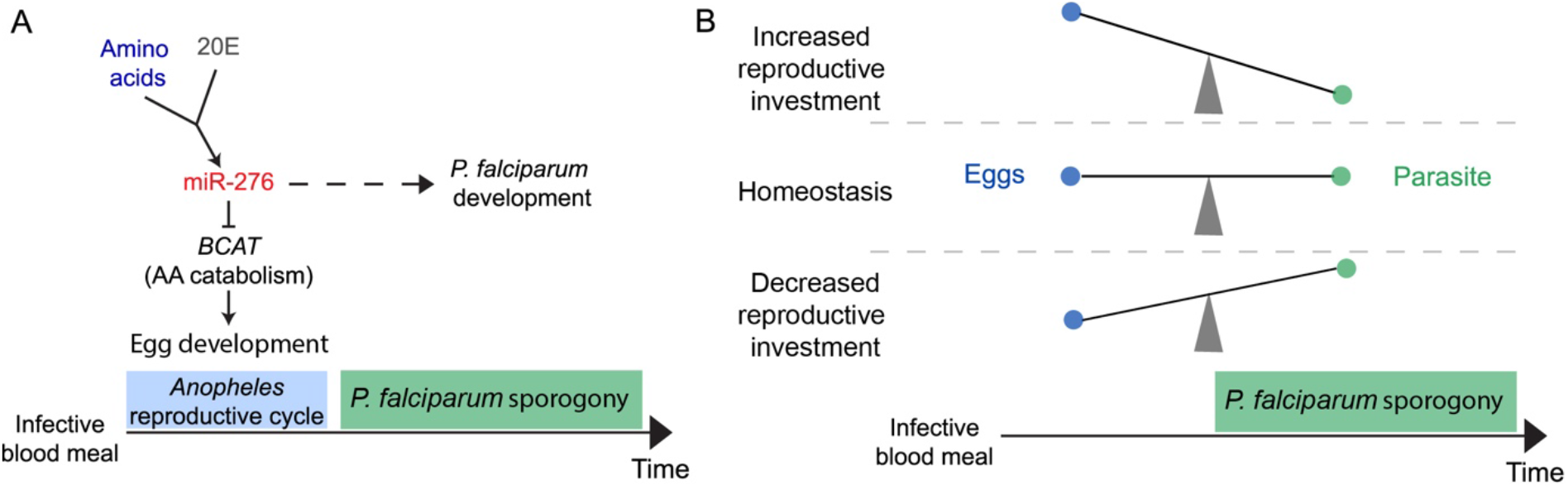
Summary of miR-276 function and proposed model of interactions between mosquito reproductive investment and *P. falciparum* development. (A) Amino acids (AA) and 20-hydroxyecdysone (20E) induce expression of miR-276 after blood feeding and, thereby, initiate the termination of the fat body AA catabolism by inhibiting the *branched chain amino acid transferase* (*BCAT*). This catabolic to anabolic switch in the fat body restricts mosquito investment into reproduction. *P. falciparum* sporogonic stages parasitize spare resources for their own development. (B) Mosquito nutritional status and the extent of the reproductive investment shape the within-vector environment which shapes *P. falciparum* sporogonic development.

Malaria parasites critically rely on mosquito hematophagy for sexual reproduction and transmission. Earlier studies focused on the trade-off between mosquito reproduction and *Plasmodium* development comparing reproductive readouts of infected versus uninfected mosquitoes (Ahmed et al., 2001; Jahan and Hurd, 1998). However, mosquito females complete egg development before massive parasite proliferation, thereby avoiding direct competition for nutrients with *Plasmodium* parasites (Costa et al., 2018). Here, we experimentally demonstrate that prolonging the reproductive cycle by inhibition of miR-276 increases mosquito reproductive investment and limits sporogonic development of *P. falciparum*. We propose that the balance between mosquito metabolic status and reproductive investment determines *Plasmodium* sporogony and transmission (Fig. 5B). Importantly, variability in reproductive investment strategies could be one of the factors underlying the differences in the vector competence observed between mosquito species that are equally exposed to *Plasmodium* infections (Briegel, 1985).

In more general terms, our study suggests that genetic or hormonal manipulations of the mosquito metabolic program and resource allocation during the reproductive cycle should impact female fertility, *Plasmodium* within-vector development and within-host virulence. We propose that the mosquito metabolic status and reproductive investment are essential components of vector competence that shape malaria transmission.

## Material and Methods

### Mosquitoes

*Anopheles coluzzii* Ngousso (*TEP1*S1*) strain was used throughout the study. Mosquitoes were maintained at 30°C 80% humidity at 12/12 h day/night cycle with a half-hour long dawn/dusk period. All mosquitoes were fed *ad libitum* with 10% sugar solution.

### Blood meal and *P. falciparum* infections

Asexual cultures (parasitaemia >2%) of *P. falciparum* NF54 (kindly provided by Prof. R. Sauerwein, RUMC, The Netherlands) were harvested by centrifugation for 5 min at 1,500 rpm and diluted with fresh red blood cells (RBS) (Haema) to 1% total parasitaemia at 4% hematocrit. Asexual and gametocyte cultures were incubated at 37°C with 3% O_2_ and 4% CO_2_. Gametocyte medium was changed daily for 15-16 days on heated plates to reduce temperature drop. On day 14 after seeding, gametocytaemia was gauged by Giemsa-stained smears, and parasite exflagellation rates were evaluated by microscopy.

Mosquitoes were fed using an artificial feeder system (Lensen et al., 1996, 1999) with diluted gametocyte cultures 15-16 days after seeding at a final gametocytaemia of 0.15 - 0.22%. The feeding system was prepared by covering the bottom of the feeders with a stretched parafilm. The feeder was heated to 37°C by water flow. Mosquitoes were fed for 15 min and unfed mosquitoes were removed. Only fully engorged females were kept for further analyses at 26°C 80% humidity.

### Parasite quantification

For *P. falciparum* oocyst counts, infected mosquitoes were killed in 70% ethanol on day 11 after infection. Mosquitoes were washed twice in PBS, midguts were dissected in 1% mercurochrome in water and incubated for 10 min at room temperature. Oocysts were counted under a stereoscope. For oocyst size, three representative images of midgut oocysts were taken per midgut by light microscope at 40x magnification and oocysts size was gauged by Fiji Image. The sporogonic index was calculated by multiplying the median number of oocysts and the mean oocyst size per experiment.

The numbers of *P. falciparum* sporozoites in the salivary glands were evaluated 14 days post infection. At least 30 mosquitoes per condition were killed in 70% ethanol, washed with PBS and the salivary glands were dissected in RPMI medium (Gibco) 3% bovine serum albumin (Sigma Aldrich). The salivary glands were homogenized with a glass plunger to release the sporozoites. The homogenate was filtered twice with cell strainers (100 μm and 40 μm) into glass vials and diluted at 1:10, 1:20 and 1:50. The sporozoites were counted using a hemocytometer under a light microscope. Sample sizes per experiment are provided in Table S4.

### 20-hydroxyecdysone (20E) quantification

20E levels were quantified using the 20E EIA kit (Cayman Chemical). Nine females per time point were collected into the microtubes, snap frozen and kept at −80°C. On the day of analysis, all samples were homogenized in 500 μl methanol using steal beads and a Qiagen tissue lyser at 50 rpm for 10 min. Samples were centrifuged at 15,000 rpm for 1 min and the supernatant (420 μl) was transferred to a clean tube. The methanol was evaporated in a speedvac. The pellets were dissolved in 230 μl of EIA buffer (Cayman Chemical). A 1:3 dilution series of 20E starting from 361 ng/ml served as a standard curve. Measurements were performed according to manufacturer’s instructions.

### *Ex vivo* fat body tissue culture

*Ex vivo* fat body culture was performed as previously described (Chung et al., 2017). In brief, three fat bodies of 3-4-day-old females were dissected and incubated with: (i) medium, (ii) 20E, (iii) amino acids (AAs); and (iv) 20E (Sigma) and AAs. 20E was dissolved in ethanol. The fat body cultures were incubated for 6 h at 27 °C. RNA was isolated by RNAzol for quantification of miRNA/mRNA expression.

### Expression analysis

#### Sample collection

For miRNA and mRNA expression kinetics, ten sugar-fed or blood-fed females were dissected on ice at different time points after a blood meal. The abdominal carcasses (the fat body) were immediately homogenized in RNAzol (Sigma Aldrich) and kept at −80°C for RNA isolation. Total RNA was isolated by RNAzol according to the manufacturer’s recommendations including the purification step with 4-Bromoanizol. The total RNA yield was measured with a Qubit (Thermo Fisher Scientific).

#### miRNA expression analysis

miRNA expression was quantified using the miScript PCR System (Qiagen) that allows real-time PCR quantification of mRNA and miRNA expression from the same reverse transcription reaction mix. The cDNA levels were further measured by Quantitect SYBR Green PCR kit (Qiagen). Forward miR-276 primer was obtained from miScript primer assay and reverse primer was purchased as previously described (Bryant et al., 2010). Expression data was calculated by the relative standard curve method. Relative quantities of miRNA expression were normalized to the gene encoding the ribosomal protein S7 (*RPS7*).

#### mRNA expression analysis

For target identification, RNAs were reverse transcribed using the RevertAid H Minus First Strand cDNA synthesis kit (Thermo Fisher Scientific) and analyzed by RT-qPCR using the SYBR Green PCR mix (Thermo Fisher Scientific). Expression data was calculated using relative standard curve method. Relative quantities of miRNA expression were normalized to the gene encoding ribosomal protein S7 (*RPS7*).

### Computational miRNA target prediction

miRNA targets were predicted *in silico* using three independent algorithms: miRANDA (John et al., 2004), RNAhybrid (Kruger and Rehmsmeier, 2006) and microTAR (Thadani and Tammi, 2006).

### miRNA inhibition

Antagomirs (anti-miR-276-5p and a scrambled version of the same antagomir that has no target in *A. gambiae* genome) were designed using the RNA module for custom single-stranded RNA synthesis (Dharmacon) as RNA anti-sense oligos with 2’Omethylated bases, a phosphorothioate backbone at the first two and last four nucleotides and a 3’cholesterol (Table S3). For miRNA inhibition, mosquitoes were anesthetized with CO2 and microinjected with 207 nl of 200 μM antagomir (41 pmol/female) at 12-18 h post eclosion using the Drummond NanoJect II (Drummond Scientific). Mosquitoes were left for four days to recover before a blood feeding or *P. falciparum* infection.

### Egg laying and hatching rates

Female and male mosquitoes were maintained in the same cage. For fertility assays, individual females were gently transferred into single cups with egg dishes (wet Whatman paper) on day three after blood feeding and allowed to oviposit for two nights. Eggs were counted by stereoscope. For fecundity assays, females were kept together after a blood feeding and allowed to lay eggs on a common egg dish. After egg laying, the egg dish was kept in water for one day at 26°C 80% humidity. Larval hatching rates were gauged by counting the number of open egg shells (at least 100 eggs) using a stereoscope.

### Dual luciferase assay

*In vitro* target validation was performed using *Drosophila* S2 cells (Invitrogen). The cells were kept in Schneider *Drosophila* medium supplemented with 10% heat-inactivated FBS (Gibco) and 1% Penicillin Streptomycin (vol/vol) (Thermo Fisher Scientific) at 25°C. Three luciferase constructs were examined: (1) a positive control containing three sites reverse complementary to miR-276 with four nucleotide linkers between each miR-276 binding site; (2) *A. gambiae BCAT* 3’UTR and (3) *A. gambiae BCAT* 3’UTR with a scrambled miR-276 binding site. All constructs were separately inserted into the multiple cloning region located downstream of the Renilla translational stop codon within the psiCheck-2 vector (Promega). psiCheck-2 reporters (100 ng) and a synthetic *A. gambiae* miR-276-5p miScript miRNA Mimic (100 ng, Qiagen) at a final concentration of 100 nm were co-transfected into *Drosophila* S2 cells using FuGENE HD transfection reagent (Promega). Cells transfected only with psiCheck-2 reporters were used as a “no miRNA mimic” control. Dual Luciferase Reporter Assay was performed 48 h post transfection using the Dual Luciferase Reporter Assay System (Promega). Firefly luciferase in the psiCheck-2 Vector was used for normalization of the Renilla luciferase expression. Measurements were made in triplicates, and transfections were repeated three times independently.

### Untargeted metabolomics

#### Sample collection

Female mosquitoes (1-day-old) were injected with anti-miR-276-5p or scrambled antagomirs as described above. Four days later, mosquitoes were infected with *P. falciparum*. At 10, 38 and 48 hpb, ten females per treatment were snap-frozen in liquid nitrogen. The samples were stored at −80°C. Additionally, at 38 and 48 hpb, abdominal carcasses (the fat body) of ten female mosquitoes per group were collected for miRNA target analysis by RT-qPCR.

#### Sample preparation

The sample preparation was performed according to MetaSysX standard procedure, a modified protocol from Giavalisco *et al* (Giavalisco et al., 2009). Whole bodies of snap-frozen mosquitoes were used for untargeted metabolomics carried out by MetaSysX, Potsdam, Germany.

#### LC-MS measurements (hydrophilic and lipophilic analytes)

The samples were measured with a Waters ACQUITY Reversed Phase Ultra Performance Liquid Chromatography (RP-UPLC) coupled to a Thermo-Fisher Q-Exactive mass spectrometer. C8 and C18 columns were used for the lipophilic and the hydrophilic measurements, respectively. Chromatograms were recorded in Full Scan MS mode (Mass Range [100-1500]).

#### LC-MS data processing (hydrophilic and lipophilic analytes)

Extraction of the LC-MS data was performed with the software REFINER MS^®^ 7.5 (GeneData, http://www.genedata.com). Alignment and filtration of the LC-MS data were completed using in-house software. After extraction of the peak list from the chromatograms, the data was processed, aligned and filtered. The alignment of the extracted data from each chromatogram was performed as follows. The samples were split into 6 groups. In order to be selected, a feature had to be present in at least 75% of replicates of at least one of the groups. At this stage, an average RT and an average *m/z* values are given to the features. The alignment was performed for each type of measurements independently (polar phase positive mode, polar phase negative mode, organic phase positive mode and organic phase negative mode). The alignment of the data was followed by the application of various filters to refine the dataset; among them the removal of isotopic peaks, the removal of in-source fragments of the analytes (due to the ionization method) and the removal of additional lower intense adduct of the same analyte.

#### LC-MS data annotation (hydrophilic and lipophilic analytes)

The annotation of the content of the sample was accomplished by matching the extracted data from the chromatograms with the library of reference compounds in terms of accurate mass and retention time.

#### MS/MS lipid annotation

Chromatograms were recorded in dd-MS2 Top 3 mode (Data Dependent tandem mass spectrometry) with the following settings: Full Scan MS mode (Mass Range [100–1500]), NCE 25 (Normalized Collision Energy). Acyl composition of di- and triacylglycerols was established from the [M+H]+ precursor ion fragmentation with detection of [Acyl+NH3] neutral losses in positive ion mode with further combinatorial calculation of the acyl composition. Acyl composition of phosphoglycerolipids was established from the detection of [Acyl-H]-fragments of the corresponding precursors in negative ion mode. Acyl composition of sphingolipids was established from the fragmentation pattern of [M+H]+ precursor ion in positive ionization mode.

#### GC-MS measurements

The samples were measured on an *Agilent* Technologies GC coupled to a *Leco Pegasus HT* mass spectrometer which consists of an EI ionization source and a TOF mass analyzer. Column: 30 m DB35; starting temp: 85°C for 2 min; gradient: 15°C per min up to 360°C.

#### GC-MS data processing and annotation

NetCDF files that were exported from the Leco Pegasus software were imported into the “R” Bioconductor package TargetSearch to transform retention time to retention index (RI), to align the chromatograms, to extract the peaks, and to annotate them by comparing the spectra and the RI to the Fiehn Library and to a user created library. Annotation of peaks was manually confirmed in Leco Pegasus. Analytes were quantified using a unique mass. Metabolites with a RT and a mass spectrum that did not have a match in the database were labelled as unknown.

#### Data filtering

The GC- and LC-MS datasets were normalized to the sample median intensity. For heatmap visualization, features with more than 50% missing values were removed and remaining missing data was replaced with the smallest value of the detected metabolite. Furthermore, the remaining dataset was subjected to interquartile range filtering, which reduced the number of features from 4716 to 2426. The data was then log2 transformed and scaled by mean-centering and by division by the square root of each feature. The raw GC-MS and LC-MS dataset is attached as supplementary material (File S1).

## Acknowledgment

The authors wish to thank Liane Spohr for mosquito breeding, Manuela Andres and Daniel Eyermann for *P. falciparum* cultures. Finally, we are grateful to all members of the vector biology unit for fruitful discussion and to Dr. Guilia Costa and Dr. Paola Carrillo-Bustamante for constructive comments on the manuscript.

## Supplementary Information

**Figure S1.**
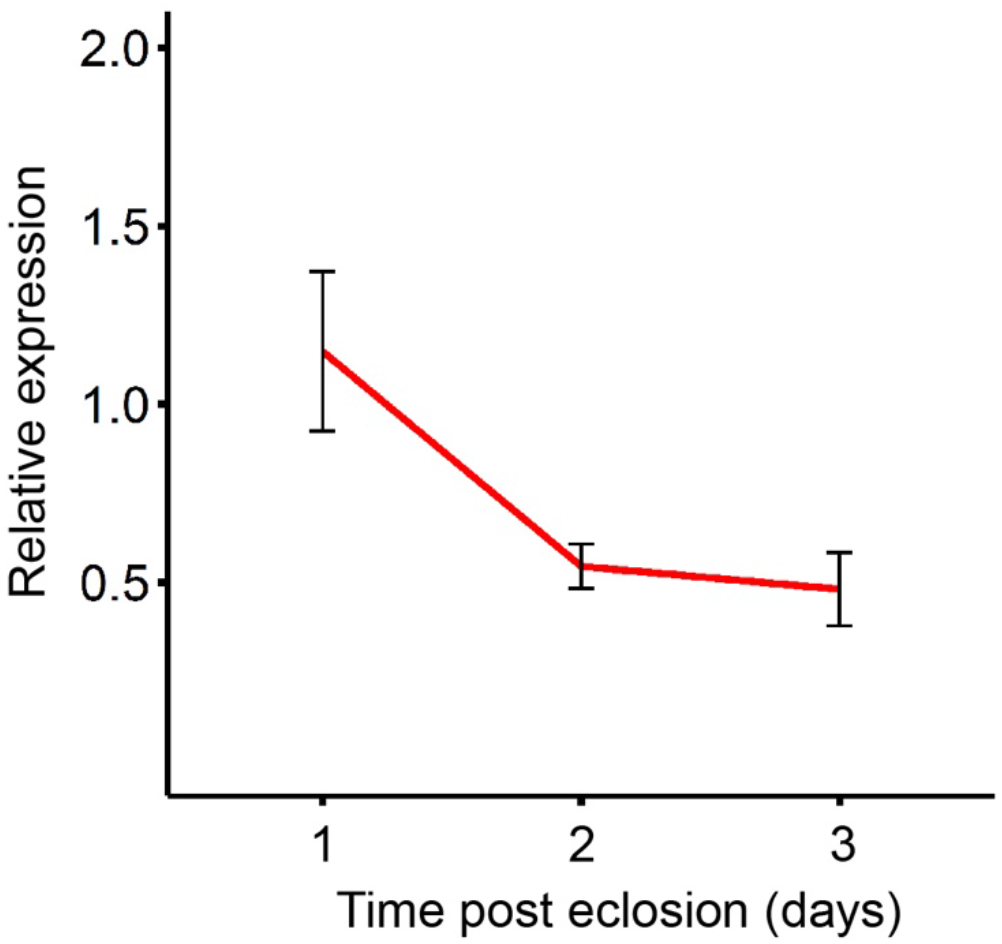
Expression of miR-276 in the fat body of mosquito females post eclosion. miR-276 expression in the fat body of mosquito females (n=10) at day 1, 2 and 3 post eclosion. Expression levels were normalized using the ribosomal protein *RPS7* gene. Data shown as mean +/− SEM (N=3) (n = number of mosquitoes pooled for each independent experiment; N = number of independent experiments).

**Figure S2.**
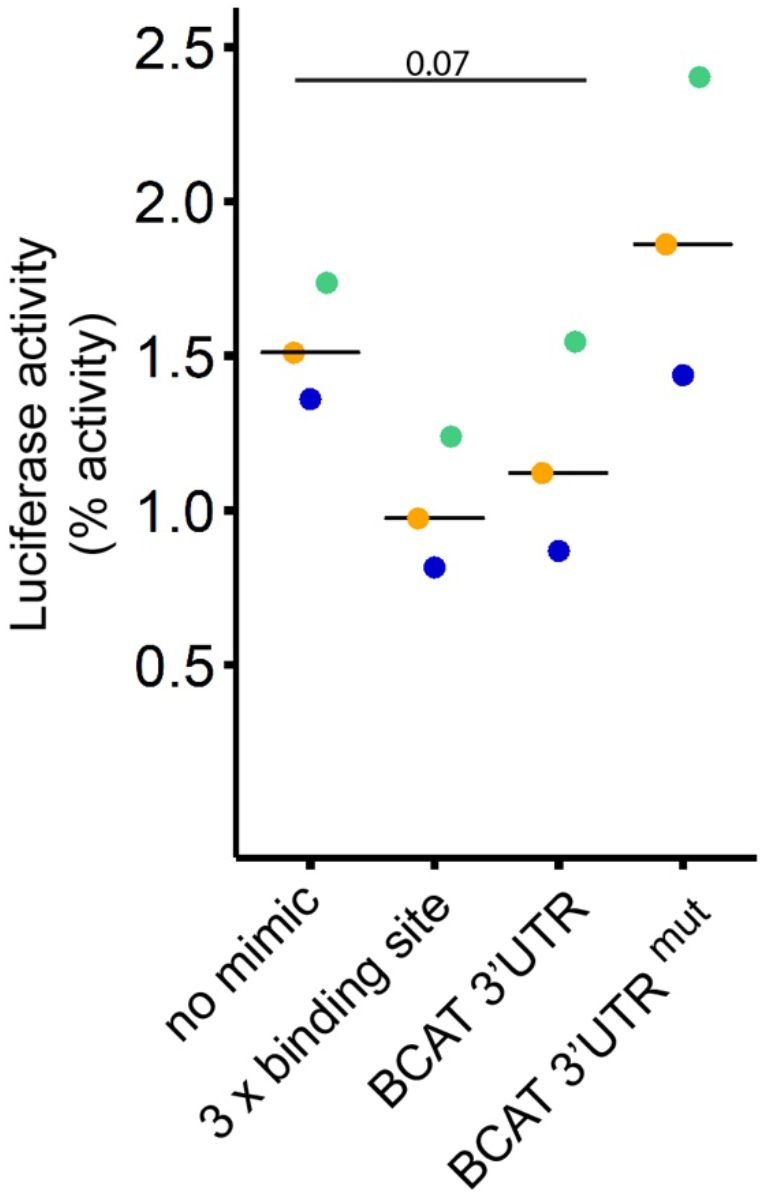
miR-276 directly targets the 3’-UTR of *BCAT*. Dual luciferase reporter assay *in vitro* using *Drosophila* S2 cells. The reporter plasmid containing three miR-276 binding sites (3 x binding site) served as a positive control. Reporter activity was induced in the absence of miR-276 mimic construct (no mimic) by the endogenous to S2 cells activity of *Drosophila* miR-276. Significant differences examined by one-way ANOVA (n=3, N=3) are shown (n = number of technical replicates within experiment; N = number of independent experiments).

**Figure S3:**
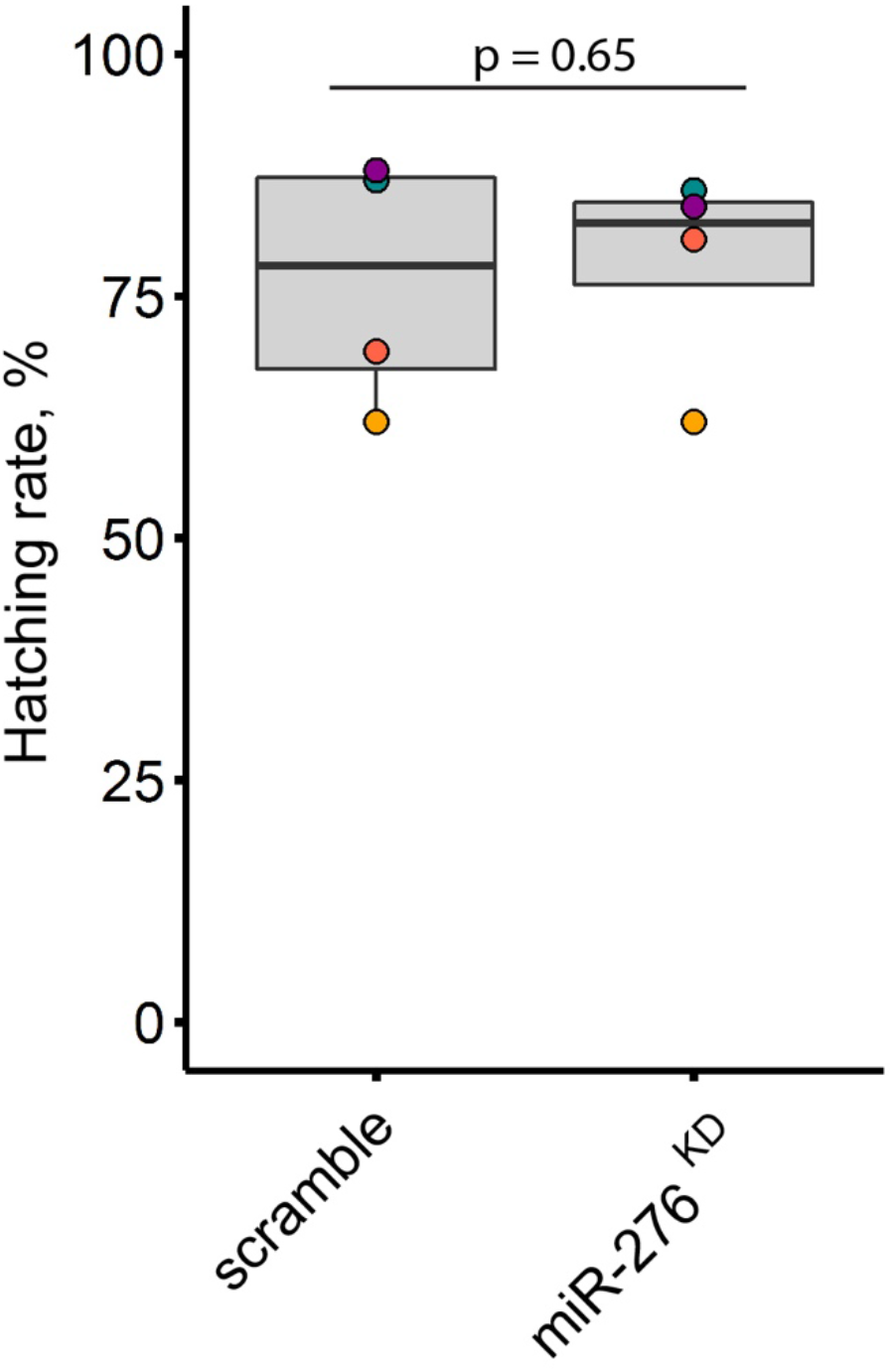
miR-276 silencing does not affect larval hatching rates. Larval hatching rates of eggs laid by females injected with anti-miR-276 (miR-276^KD^) or scrambled antagomir (scramble). Boxplots show the median with first and third quartile, whiskers depict min and max values. Each dot color represents one independent experiment with at least 100 eggs (N = 4, N = number of independent experiments).

**Table S1:** miRNA target prediction. The miRANDA, RNAhybrid and MicroTar algorithm were used to predict miR-276-5p targets.

| Gene ID | Annotation | miRanda | RNAhybrid | MicroTar |
| --- | --- | --- | --- | --- |
| AGAP010034 | Mediator of RNA polymerase II transcription subunit 6 | YES | YES | YES |
| AGAP009633 | 3-oxoacyl-(acyl-carrier protein) reductase fabG2 | YES | YES | YES |
| AGAP006009 | Cuticular protein RR-1 family 30 | YES | YES | YES |
| AGAP012081 | F-type H<sup>+</sup>-transporting ATPase subunit beta | YES | YES | YES |
| AGAP012726 | Stabilizer of axonemal microtubules 1/2 | YES | YES | YES |
| AGAP010983 | BAG family molecular chaperone regulator 2 | YES | YES | YES |
| AGAP001250 | Eupolytin | YES | YES | YES |
| AGAP009882 | Heat shock factor-binding protein 1 | YES | YES | YES |
| AGAP002227 | Pyridoxamine 5'-phosphate oxidase | YES | YES | YES |
| AGAP005818 | Nurim homolog | YES | YES | YES |
| AGAP011129 |  | YES | YES | YES |
| AGAP013228 | Heat shock protein 67B2 | YES | YES | YES |
| AGAP006549 |  | YES | YES | YES |
| AGAP003108 | Fibroblast growth factor receptor 2 | YES | YES | YES |
| AGAP010163 | 60S ribosomal protein L38 | YES | YES | YES |
| AGAP002603 | Elongation factor 1 alpha-like protein | YES | YES | YES |
| AGAP004490 |  | YES | YES | YES |
| AGAP012910 | GPI inositol-deacylase | YES | YES | YES |
| AGAP008478 | Cuticular protein CPLCW family | YES | YES | YES |
| AGAP006184 |  | YES | YES | YES |
| AGAP001801 | DTW domain-containing protein 2 | YES | YES | YES |
| **AGAP000011** | **Branched-chain-amino-acid aminotransferase 2** | **YES** | **YES** | **YES** |
| AGAP006355 | Transcription initiation factor TFIIE subunit alpha | YES | YES | YES |
| AGAP008194 |  | YES | YES | YES |
| AGAP002927 | High affinity cGMP-specific 3',5'-cyclic phosphodiesterase 9 | YES | YES | YES |
| AGAP005003 | Melanotransferrin | YES | YES | YES |
| AGAP006532 | UDP-glucose 6-dehydrogenase | YES | YES | YES |
| AGAP002053 |  | YES | YES | YES |
| AGAP013070 |  | YES | YES | YES |
| AGAP009935 |  | YES | YES | YES |
| AGAP000768 | Septin 4 | YES | YES | YES |
| AGAP004952 | CBF1 interacting corepressor | YES | YES | YES |
| AGAP010824 |  | YES | YES | YES |
| AGAP007872 | Mps one binder kinase activator-like 3 | YES | YES | YES |
| AGAP006586 | Serine protease inhibitor-like superfamily | YES | YES | YES |
| AGAP013323 |  | YES | YES | YES |
| AGAP000882 | 17-beta-hydroxysteroid dehydrogenase type 6 precursor | YES | YES | YES |
| AGAP002439 | Cytochrome c oxidase assembly protein COX14 | YES | YES | YES |
| AGAP000969 | Topomodulin | YES | YES | YES |
| AGAP000342 | Inhibin, beta | YES | YES | YES |
| AGAP003356 | KH domain-containing, RNA-binding, signal transduction-associated protein 3 | YES | YES | YES |
| AGAP007990 | Glucosyl/glucuronosyl transferases | YES | YES | YES |
| AGAP003810 | Kune | YES | YES | YES |
| AGAP005611 |  | YES | YES | YES |
| AGAP007476 | TP53 regulating kinase | YES | YES | YES |
| AGAP002113 | Cytochrome b5 | YES | YES | YES |
| AGAP002067 | Cell cycle progression | YES | YES | YES |
| AGAP012847 | Autophagy-related protein 12 | NO | YES | YES |
| AGAP007851 |  | NO | YES | YES |
| AGAP001630 | Triosephosphate isomerase | NO | YES | YES |
| AGAP006678 | Cyclin T | NO | YES | YES |
| AGAP002198 | Glycine N-methyltransferase | NO | YES | YES |
| AGAP002239 | Diamine acetyltransferase 1 | NO | YES | YES |
| AGAP008499 | Mitochondrial transcription factor A | NO | YES | YES |
| AGAP006611 | TRAF-interacting protein | NO | YES | YES |
| AGAP004788 | Up-regulated during skeletal muscle growth 5 homolog | NO | YES | YES |
| AGAP006593 | Dual specificity phosphatase | NO | YES | YES |
| AGAP000607 | SG1-like salivary protein | NO | YES | YES |
| AGAP006479 | Myosin V | NO | YES | YES |
| AGAP007997 |  | NO | YES | YES |
| AGAP000080 |  | NO | YES | YES |
| AGAP003494 | Sugar transporter ERD6-like 7 | NO | YES | YES |
| AGAP010228 | 3-methylcrotonyl-CoA carboxylase alpha subunit | NO | YES | YES |
| AGAP001277 |  | NO | YES | YES |
| AGAP003019 | Cleavage stimulation factor subunit 3 | NO | YES | YES |
| AGAP005031 | E3 SUMO-protein ligase PIAS2 | NO | YES | YES |
| AGAP012654 |  | NO | YES | YES |
| AGAP005079 | Guanine nucleotide-binding protein G(q) alpha subunit | NO | YES | YES |
| AGAP008779 |  | NO | YES | YES |
| AGAP004668 | Coiled-coil domain-containing protein 12 | NO | YES | YES |
| AGAP001161 | Long wavelength sensitive opsin | NO | YES | YES |
| AGAP009579 | Dihydropyridine-sensitive l-type calcium channel | NO | YES | YES |
| AGAP001894 | Arrestin domain-containing protein 3 | NO | YES | YES |
| AGAP000545 |  | NO | YES | YES |
| AGAP007411 | C-type lectin (CTL) - mannose binding | NO | YES | YES |
| AGAP002429 | Cytochrome P450, shade | NO | YES | YES |
| AGAP010782 | Ral guanine nucleotide exchange factor 2 | NO | YES | YES |
| AGAP002256 | RNA-binding protein 7 | NO | YES | YES |
| AGAP013773 |  | NO | YES | YES |
| AGAP003109 | Transcriptional adapter 2-beta | NO | YES | YES |
| AGAP005218 | Zinc finger Ran-binding domain-containing protein 2 | NO | YES | YES |
| AGAP012787 |  | NO | YES | YES |
| AGAP009687 | U6 snRNA-associated Sm-like protein LSm4 | NO | YES | YES |
| AGAP010098 | Cuticular protein RR-2 family 83 | NO | YES | YES |
| AGAP002677 | Coiled-coil domain-containing protein lobo homolog | NO | YES | YES |
| AGAP000500 | NADPH cytochrome P450 reductase | NO | YES | YES |
| AGAP003482 |  | NO | YES | YES |
| AGAP001215 | Tetratricopeptide repeat protein 39C | NO | YES | YES |
| AGAP011838 |  | NO | YES | YES |
| AGAP006806 | Mitochondrial thiamine pyrophosphate carrier | NO | YES | YES |
| AGAP003337 |  | NO | YES | YES |
| AGAP002216 | Cation transport regulator-like protein 2 | NO | YES | YES |
| AGAP008018 | Cytochrome P450 | NO | YES | YES |
| AGAP006505 |  | NO | YES | YES |
| AGAP004992 | Microsomal prostaglandin-E synthase 2 | NO | YES | YES |
| AGAP002395 | 60S ribosomal protein L10-2 | NO | YES | YES |
| AGAP007418 | Proteasome inhibitor subunit 1 (PI31) | NO | YES | YES |
| AGAP002078 | 14 kDa phosphohistidine phosphatase | NO | YES | YES |
| AGAP011680 | Facilitated glucose transporter (solute carrier family 2) | NO | YES | YES |
| AGAP005952 |  | NO | YES | YES |
| AGAP013283 |  | NO | YES | YES |
| AGAP007296 | Autophagy related gene | NO | YES | YES |
| AGAP012030 | 39S ribosomal protein L21, mitochondrial | NO | YES | YES |
| AGAP002587 |  | NO | YES | YES |
| AGAP011753 |  | NO | YES | YES |
| AGAP013210 | Down syndrome cell adhesion molecule-like protein 1 | NO | YES | YES |
| AGAP006253 | Cysteine-rich venom protein | NO | YES | YES |
| AGAP012895 | Histone H2A | YES | NO | YES |
| AGAP006937 | Ubiquitin-conjugating enzyme E2 variant | YES | NO | YES |
| AGAP008227 | Trehalose 6-phosphate synthase/phosphatase | YES | NO | YES |
| AGAP000141 |  | YES | NO | YES |
| AGAP011424 | 40S ribosomal protein S16 | YES | NO | YES |
| AGAP006914 | Fibrinogen-related protein 1 | YES | NO | YES |
| AGAP007835 |  | YES | NO | YES |
| AGAP005744 | Leucine-rich immune protein | YES | NO | YES |
| AGAP005860 | Phosphoglucomutase | YES | NO | YES |
| AGAP011235 |  | YES | NO | YES |
| AGAP006991 |  | YES | NO | YES |
| AGAP009852 | Mitochondrial fission 1 protein | YES | NO | YES |
| AGAP001139 | 39S ribosomal protein L19, mitochondrial | YES | NO | YES |
| AGAP003790 | Annexin B9 | YES | NO | YES |
| AGAP001418 |  | YES | NO | YES |
| AGAP007650 | Growth arrest and DNA-damage-inducible protei | YES | NO | YES |
| AGAP000109 | Cytochrome c oxidase subunit VIIa | YES | NO | YES |
| AGAP008862 | Aubergine | YES | NO | YES |
| AGAP003545 | Protein oskar | YES | NO | YES |
| AGAP001083 |  | YES | NO | YES |
| AGAP009777 | Mothers against decapentaplegic homolog 2/3 | YES | NO | YES |
| AGAP003390 | Cuticular protein RR-2 family 124 | YES | NO | YES |
| AGAP002499 | Methylmalonate-semialdehyde dehydrogenase (acylating), mitochondrial | YES | NO | YES |
| AGAP001711 | NADH dehydrogenase (ubiquinone) Fe-S protein 8 | YES | NO | YES |
| AGAP007940 | Reticulon-like protein | YES | NO | YES |
| AGAP001368 |  | YES | NO | YES |
| AGAP007952 | Elongation factor 1 alpha-like protein | YES | NO | YES |
| AGAP003939 |  | YES | NO | YES |
| AGAP008648 |  | YES | NO | YES |
| AGAP005776 | Pigment dispersing hormone | YES | NO | YES |
| AGAP012106 | F-type H<sup>+</sup>-transporting ATPase subunit beta | YES | NO | YES |
| AGAP003440 |  | YES | NO | YES |
| AGAP008974 | Protein cornichon homolog | YES | NO | YES |
| AGAP006933 |  | YES | NO | YES |
| AGAP002159 |  | YES | NO | YES |
| AGAP006747 | IMD pathway signalling NF-kappaB Relish-like transcription factor | YES | NO | YES |
| AGAP006829 | Cuticular protein RR-1 family 59 | YES | NO | YES |
| AGAP005324 | Ubiquitin-conjugating enzyme E2 W | YES | NO | YES |
| AGAP003596 |  | YES | NO | YES |
| AGAP002114 | Cell division cycle 20-like protein 1, cofactor of APC complex | YES | NO | YES |
| AGAP008292 | Trypsin 4 | YES | NO | YES |
| AGAP010217 | Protein disulfide-isomerase | YES | NO | YES |
| AGAP012207 |  | YES | NO | YES |
| AGAP000774 | PH and SEC7 domain-containing protein | YES | NO | YES |
| AGAP007738 |  | YES | NO | YES |
| AGAP000044 | Thioredoxin domain-containing protein | YES | NO | YES |
| AGAP000983 |  | YES | NO | YES |
| AGAP007928 | Turtle protein, isoform | YES | NO | YES |
| AGAP011369 | Gelsolin | YES | NO | YES |
| AGAP003136 | 2-oxoisovalerate dehydrogenase E1 component, alpha subunit | YES | NO | YES |
| AGAP000348 |  | YES | NO | YES |
| AGAP011323 |  | YES | NO | YES |
| AGAP011945 |  | YES | NO | YES |
| AGAP002308 | Pyrroline-5-carboxylate reductase | YES | NO | YES |
| AGAP009464 | ATP-binding cassette transporter (ABC transporter) family G member 10 | YES | NO | YES |
| AGAP005781 | Glycine C-acetyltransferase | YES | NO | YES |
| AGAP012186 | Lipoyltransferase 1 | YES | NO | YES |
| AGAP003874 | Uridine kinase | YES | NO | YES |
| AGAP003580 |  | YES | NO | YES |
| AGAP007649 |  | YES | NO | YES |
| AGAP002982 | E3 SUMO-protein ligase RanBP2 | YES | NO | YES |
| AGAP005369 |  | YES | NO | YES |
| AGAP000039 | Gamma-aminobutyric acid receptor subunit alpha | YES | NO | YES |
| AGAP010365 |  | YES | YES | NO |
| AGAP012177 | Aveugle | YES | YES | NO |
| AGAP000692 | Cecropin-C | YES | YES | NO |
| AGAP008578 | ATP-dependent RNA helicase DDX4 | YES | YES | NO |
| AGAP002047 | Atlastin | YES | YES | NO |
| AGAP002063 | Nucleolar protein 56 | YES | YES | NO |
| AGAP004513 | 4-hydroxybenzoate hexaprenyltransferase | YES | YES | NO |
| AGAP000199 | Phospholipase D3/4 | YES | YES | NO |
| AGAP007133 |  | YES | YES | NO |
| AGAP005612 |  | YES | YES | NO |
| AGAP006070 | Prefoldin 2 | YES | YES | NO |
| AGAP002244 |  | YES | YES | NO |
| AGAP005525 |  | YES | YES | NO |
| AGAP012391 |  | YES | YES | NO |
| AGAP003331 | Enolase-phosphatase E1 | YES | YES | NO |
| AGAP001458 | Mono-ADP-ribosyltransferase sirtuin 6 | YES | YES | NO |
| AGAP004437 | Glycerol-3-phosphate dehydrogenase | YES | YES | NO |
| AGAP009979 | gGlycoprotein-N-acetylgalactosamine 3-beta-galactosyltransferase | YES | YES | NO |
| AGAP001980 |  | YES | YES | NO |
| AGAP010840 |  | YES | YES | NO |
| AGAP008813 | FMS-like tyrosine kinase 1 | YES | YES | NO |
| AGAP003768 | 40S ribosomal protein S2 | YES | YES | NO |
| AGAP010880 |  | YES | YES | NO |
| AGAP001308 | Coiled-coil domain-containing protein 64 | YES | YES | NO |
| AGAP007119 | Sideroflexin 1,2,3 | YES | YES | NO |
| AGAP001889 |  | YES | YES | NO |
| AGAP004280 | Groucho protein | YES | YES | NO |
| AGAP010464 | NADH dehydrogenase (ubiquinone) 1 alpha/beta subcomplex 1 | YES | YES | NO |
| AGAP013250 |  | YES | YES | NO |
| AGAP001779 |  | YES | YES | NO |
| AGAP008445 | Cuticular protein CPLCG family (CPLCG2) | YES | YES | NO |
| AGAP007889 | UDP-N-acetylglucosamine pyrophosphorylase | YES | YES | NO |
| AGAP012972 |  | YES | YES | NO |
| AGAP009105 | Serine/threonine-protein phosphatase 2A 65 kDa regulatory subunit A beta isoform isoform c | YES | YES | NO |
| AGAP005245 |  | YES | YES | NO |
| AGAP000448 | Solute carrier family 25 (mitochondrial ornithine transporter) member 2/15 | YES | YES | NO |
| AGAP002203 |  | YES | YES | NO |
| AGAP001306 | Actin related protein 2/3 complex, subunit 4 | YES | YES | NO |
| AGAP005210 | Translation initiation factor eIF-2B subunit gamma | YES | YES | NO |
| AGAP009361 |  | YES | YES | NO |
| AGAP003132 |  | YES | YES | NO |
| AGAP002933 | 2-oxoglutarate and iron-dependent oxygenase domain-containing protein 1 | YES | YES | NO |
| AGAP003746 |  | YES | YES | NO |
| AGAP004283 | AP-1 complex subunit sigma 1/2 | YES | YES | NO |
| AGAP004825 |  | YES | YES | NO |
| AGAP001417 | Brain protein 44-like protein | YES | YES | NO |
| AGAP010672 | Succinate dehydrogenase (ubiquinone) cytochrome b560 subunit | YES | YES | NO |
| AGAP005038 |  | YES | YES | NO |
| AGAP012960 |  | YES | YES | NO |
| AGAP000801 | ilnotropic receptor GLURIIb | YES | YES | NO |
| AGAP004510 | Innexin inx2 | YES | YES | NO |
| AGAP002630 | NADH dehydrogenase (ubiquinone) 1 beta subcomplex 2 | YES | YES | NO |
| AGAP000093 |  | YES | YES | NO |
| AGAP013534 |  | YES | YES | NO |
| AGAP009434 | Valacyclovir hydrolase | YES | YES | NO |
| AGAP002556 | Odorant-binding protein 66 | YES | YES | NO |
| AGAP005471 |  | YES | YES | NO |
| AGAP009024 | RNA polymerase I-specific transcription initiation factor RRN3 | YES | YES | NO |
| AGAP001601 | Ser/Thr protein phosphatase/nucleotidase | YES | YES | NO |
| AGAP005748 |  | YES | YES | NO |
| AGAP002507 | Peroxisomal biogenesis factor 3 | YES | YES | NO |
| AGAP011026 | 5' nucleotidase, ecto | YES | YES | NO |
| AGAP002782 | Nuclear autoantigenic sperm protein | YES | YES | NO |
| AGAP000070 | Protein kinase | YES | YES | NO |
| AGAP006344 | RAG1-activating protein 1-like protein | YES | YES | NO |
| AGAP013730 |  | YES | YES | NO |
| AGAP013023 |  | YES | YES | NO |

**Table S2:**
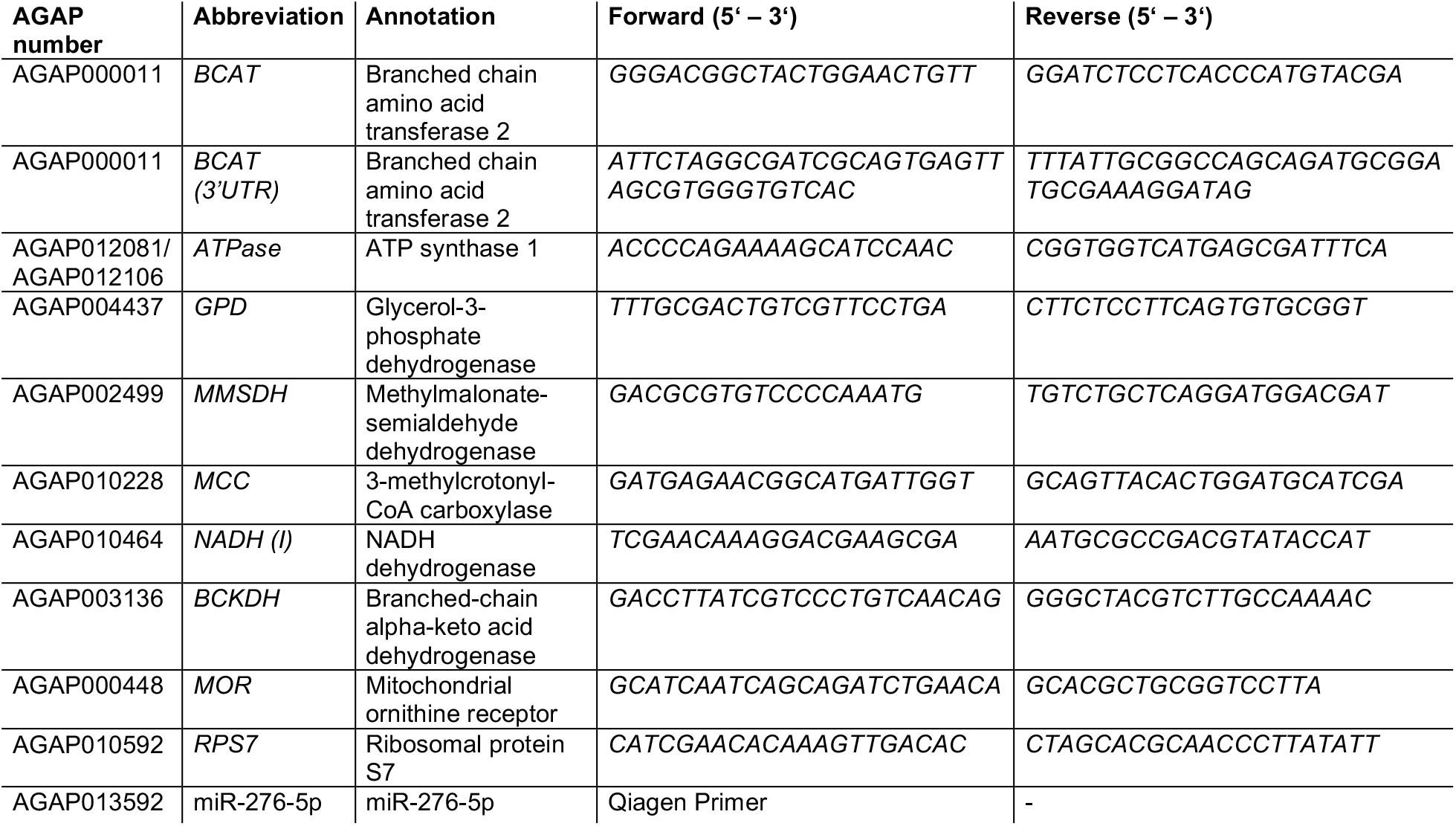
Primers used in this study.

**Table S3:** Antagomir list

| Name | Sequence |
| --- | --- |
| anti-miR-276-5p | 5'-U.*.A.*.G.G.A.A.C.U.C.U.A.U.A.C.C.U.C.*.G.*.C.*.U.*.A.mN.3'-Chl-3' |
| scramble | 5'-A.*.A.*.U.G.G.A.C.C.C.U.A.C.U.A.A.U.U.*.U.*.G.*.C.*.C.mN.3'-Chl-3' |

\* = phosphothioate backbone; mN = OCH<sub>3</sub>-group; Chl = Cholesterol

**Table S4:** Sample sizes in infection and fertility experiments

| **Experiment** | **Replicate** | **Unit of n** | **n per experiment** |
| --- | --- | --- | --- |
| Egg laying rate | Replicate 1 | mosquito | 12 (scramble);<br>9 (miR-276<sup>KD</sup>) |
|  | Replicate 2 | mosquito | 13 (scramble);<br>14 (miR-276<sup>KD</sup>) |
|  | Replicate 3 | mosquito | 17 (scramble);<br>16 (miR-276<sup>KD</sup>) |
| Oocyst number | Replicate 1 | midgut | 25 (scramble);<br>28 (miR-276<sup>KD</sup>) |
|  | Replicate 2 | midgut | 16 (scramble);<br>17 (miR-276<sup>KD</sup>) |
|  | Replicate 3 | midgut | 21 (scramble);<br>20 (miR-276<sup>KD</sup>) |
| Oocyst size | Replicate 1 | oocyst | 191 (scramble);<br>87 (miR-276<sup>KD</sup>) |
|  | Replicate 2 | oocyst | 117 (scramble);<br>131 (miR-276<sup>KD</sup>) |
|  | Replicate 3 | oocyst | 85 (scramble);<br>70 (miR-276<sup>KD</sup>) |
| Sporozoites number | Replicate 1 | mosquito salivary gland | 30 (scramble); pooled<br>30 (miR-276<sup>KD</sup>); pooled |
|  | Replicate 2 | mosquito salivary gland | 30 (scramble); pooled<br>30 (miR-276<sup>KD</sup>); pooled |
|  | Replicate 3 | mosquito salivary gland | 30 (scramble); pooled<br>30 (miR-276<sup>KD</sup>); pooled |

## References

Ahmed, a. M., Maingon, R., Romans, P., and Hurd, H. (2001). Effects of malaria infection on vitellogenesis in Anopheles gambiae during two gonotrophic cycles. Insect Molecular Biology 10, 347–356.

Arrese, E.L., and Soulages, J.L. (2010). Insect fat body: energy, metabolism, and regulation. Annual Review of Entomology 55, 207–225.

Briegel, H. (1985). Mosquito reproduction: Incomplete utilization of the blood meal protein for oögenesis. Journal of Insect Physiology 31, 15–21.

Briegel, H., Hefti, M., and DiMarco, E. (2002). Lipid metabolism during sequential gonotrophic cycles in large and small female Aedes aegypti. Journal of Insect Physiology 48, 547–554.

Bryant, B., Macdonald, W., and Raikhel, A.S. (2010). microRNA miR-275 is indispensable for blood digestion and egg development in the mosquito Aedes aegypti. Proceedings of the National Academy of Sciences of the United States of America 107, 22391–22398.

Bryant, B., Raikhel, A.S., Bambina, S., Berman, A., and Cherry, S. (2011). Programmed Autophagy in the Fat Body of Aedes aegypti Is Required to Maintain Egg Maturation Cycles. PLoS ONE 6, e25502.

Carpenter, V.K., Drake, L.L., Aguirre, S.E., Price, D.P., Rodriguez, S.D., and Hansen, I.A. (2012). SLC7 amino acid transporters of the yellow fever mosquito Aedes aegypti and their role in fat body TOR signaling and reproduction. Journal of Insect Physiology 58, 513–522.

Chung, H.-N., Rodriguez, S.D., Carpenter, V.K., Vulcan, J., Bailey, C.D., Nageswara-Rao, M., Li, Y., Attardo, G.M., and Hansen, I.A. (2017). Fat Body Organ Culture System in Aedes Aegypti, a Vector of Zika Virus. Journal of Visualized Experiments e55508–e55508.

Costa, G., Gildenhard, M., Eldering, M., Lindquist, R.L., Hauser, A.E., Sauerwein, R., Goosmann, C., Brinkmann, V., Carrillo-Bustamante, P., and Levashina, E.A. (2018). Non-competitive resource exploitation within mosquito shapes within-host malaria infectivity and virulence. Nature Communications 9, 3474.

Deitsch, K.W., Chen, J.-S., and Raikhel, A.S. (1995). Indirect control of yolk protein genes by 20-hydroxyecdysone in the fat body of the mosquito, Aedes aegypti. Insect Biochemistry and Molecular Biology 25, 449–454.

Esslinger, S.M., Schwalb, B., Helfer, S., Michalik, K.M., Witte, H., Maier, K.C., Martin, D., Michalke, B., Tresch, A., Cramer, P., et al. (2013). Drosophila miR-277 controls branched-chain amino acid catabolism and affects lifespan. RNA Biology 10, 1042–1056.

Fuchs, S., Behrends, V., Bundy, J.G., Crisanti, A., and Nolan, T. (2014a). Phenylalanine metabolism regulates reproduction and parasite melanization in the malaria mosquito. PLoS ONE 9, e84865.

Fuchs, S., Behrends, V., Bundy, J.G., Crisanti, A., and Nolan, T. (2014b). Phenylalanine Metabolism Regulates Reproduction and Parasite Melanization in the Malaria Mosquito. PLoS ONE 9, e84865.

Giavalisco, P., Köhl, K., Hummel, J., Seiwert, B., and Willmitzer, L. (2009). ^13^ C Isotope-Labeled Metabolomes Allowing for Improved Compound Annotation and Relative Quantification in Liquid Chromatography-Mass Spectrometry-based Metabolomic Research. Analytical Chemistry 81, 6546–6551.

Gonçalves, R.L.S., Machado, A.C.L., Paiva-Silva, G.O., Sorgine, M.H.F., Momoli, M.M., Oliveira, J.H.M., Vannier-Santos, M.A., Galina, A., Oliveira, P.L., and Oliveira, M.F. (2009). Blood-feeding induces reversible functional changes in flight muscle mitochondria of Aedes aegypti mosquito. PloS ONE 4, e7854.

Gulia-Nuss, M., Robertson, A.E., Brown, M.R., Strand, M.R., and Pankratz, M. (2011). Insulin-Like Peptides and the Target of Rapamycin Pathway Coordinately Regulate Blood Digestion and Egg Maturation in the Mosquito Aedes aegypti. PLoS ONE 6, e20401.

Hansen, I.A., Attardo, G.M., Park, J.-H., Peng, Q., and Raikhel, A.S. (2004). Target of rapamycin-mediated amino acid signaling in mosquito anautogeny. Proceedings of the National Academy of Sciences of the United States of America 101, 10626–10631.

Hansen, I.A., Attardo, G.M., Roy, S.G., and Raikhel, A.S. (2005). Target of rapamycin-dependent activation of S6 kinase is a central step in the transduction of nutritional signals during egg development in a mosquito. The Journal of Biological Chemistry 280, 20565–20572.

Hou, Y., Wang, X.-L., Saha, T.T., Roy, S., Zhao, B., Raikhel, A.S., and Zou, Z. (2015). Temporal Coordination of Carbohydrate Metabolism during Mosquito Reproduction. PLoS Genetics 11, e1005309.

Isoe, J., and Scaraffia, P.Y. (2013). Urea Synthesis and Excretion in Aedes aegypti Mosquitoes Are Regulated by a Unique Cross-Talk Mechanism. PLoS ONE 8, e65393.

Isoe, J., Petchampai, N., Isoe, Y.E., Co, K., Mazzalupo, S., and Scaraffia, P.Y. (2017). Xanthine dehydrogenase-1 silencing in *Aedes aegypti* mosquitoes promotes a blood feeding–induced adulticidal activity. The FASEB Journal 31, 2276–2286.

Jahan, N., and Hurd, H. (1998). Effect of Plasmodium yoelii nigeriensis (Haemosporidia: Plasmodiidae) on Anopheles stephensi (Diptera: Culicidae) vitellogenesis. Journal of Medical Entomology 35, 956–961.

John, B., Enright, A.J., Aravin, A., Tuschl, T., Sander, C., and Marks, D.S. (2004). Human MicroRNA Targets. PLoS Biology 2, e363.

Kruger, J., and Rehmsmeier, M. (2006). RNAhybrid: microRNA target prediction easy, fast and flexible. Nucleic Acids Research 34, W451–W454.

Lampe, L., and Levashina, E.A. (2018). MicroRNA Tissue Atlas of the Malaria Mosquito Anopheles gambiae. G3 (Bethesda, Md.) 8, 185–193.

Lensen, A., van Druten, J., Bolmer, M., van Gemert, G., Eling, W., and Sauerwein, R. (1996). Measurement by membrane feeding of reduction in Plasmodium falciparum transmission induced by endemic sera. Transactions of the Royal Society of Tropical Medicine and Hygiene 90, 20–22.

Lensen, A., Bril, A., Van de Vegte, M., Gemert, G.J. van, Eling, W., and Sauerwein, R. (1999). Infectivity of cultured Plasmodium falciparum gametocytes to mosquitoes. Experimental Parasitology 91, 101–103.

Li, T., Zhang, Z., Kolwicz, S.C., Abell, L., Roe, N.D., Kim, M., Zhou, B., Cao, Y., Ritterhoff, J., Gu, H., et al. (2017). Defective Branched-Chain Amino Acid Catabolism Disrupts Glucose Metabolism and Sensitizes the Heart to Ischemia-Reperfusion Injury. Cell Metabolism 25, 374–385.

Liu, S., Lucas, K.J., Roy, S., Ha, J., and Raikhel, A.S. (2014). Mosquito-specific microRNA-1174 targets serine hydroxymethyltransferase to control key functions in the gut. Proceedings of the National Academy of Sciences of the United States of America 111, 14460–14465.

Lucas, K.J., Zhao, B., Roy, S., Gervaise, A.L., and Raikhel, A.S. (2015). Mosquito-specific microRNA-1890 targets the juvenile hormone-regulated serine protease JHA15 in the female mosquito gut. RNA Biology 12, 1383–1390.

Magalhaes, T., Brackney, D.E., Beier, J.C., and Foy, B.D. (2008). Silencing an Anopheles gambiae catalase and sulfhydryl oxidase increases mosquito mortality after a blood meal. Archives of Insect Biochemistry and Physiology 68, 134–143.

Mane-Padros, D., Cruz, J., Cheng, A., and Raikhel, A.S. (2012a). A critical role of the nuclear receptor HR3 in regulation of gonadotrophic cycles of the mosquito Aedes aegypti. PloS ONE 7, e45019.

Mane-Padros, D., Cruz, J., Cheng, A., Raikhel, A.S., Hansen, I., Attardo, G., Park, J., Peng, Q., Raikhel, A., Roy, S., et al. (2012b). A Critical Role of the Nuclear Receptor HR3 in Regulation of Gonadotrophic Cycles of the Mosquito Aedes aegypti. PLoS ONE 7, e45019.

Mazzalupo, S., Isoe, J., Belloni, V., and Scaraffia, P.Y. (2016). Effective disposal of nitrogen waste in blood-fed Aedes aegypti mosquitoes requires alanine aminotransferase. FASEB Journal : Official Publication of the Federation of American Societies for Experimental Biology 30, 111–120.

Moller-Jacobs, L.L., Murdock, C.C., and Thomas, M.B. (2014). Capacity of mosquitoes to transmit malaria depends on larval environment. Parasites & Vectors 7, 593.

Rono, M.K., Whitten, M.M.A., Oulad-Abdelghani, M., Levashina, E.A., and Marois, E. (2010). The major yolk protein vitellogenin interferes with the anti-plasmodium response in the malaria mosquito Anopheles gambiae. PLoS Biology 8, e1000434.

Roy, S., Saha, T.T., Johnson, L., Zhao, B., Ha, J., White, K.P., Girke, T., Zou, Z., and Raikhel, A.S. (2015). Regulation of Gene Expression Patterns in Mosquito Reproduction. PLoS Genetics 11, e1005450.

Scaraffia, P.Y., Zhang, Q., Wysocki, V.H., Isoe, J., and Wells, M.A. (2006). Analysis of whole body ammonia metabolism in Aedes aegypti using [15N]-labeled compounds and mass spectrometry. Insect Biochemistry and Molecular Biology 36, 614–622.

Scaraffia, P.Y., Zhang, Q., Thorson, K., Wysocki, V.H., and Miesfeld, R.L. (2010). Differential ammonia metabolism in Aedes aegypti fat body and midgut tissues. Journal of Insect Physiology 56, 1040–1049.

She, P., Reid, T.M., Bronson, S.K., Vary, T.C., Hajnal, A., Lynch, C.J., and Hutson, S.M. (2007). Disruption of BCATm in Mice Leads to Increased Energy Expenditure Associated with the Activation of a Futile Protein Turnover Cycle. Cell Metabolism 6, 181–194.

Short, S.M., Mongodin, E.F., MacLeod, H.J., Talyuli, O.A.C., and Dimopoulos, G. (2017). Amino acid metabolic signaling influences Aedes aegypti midgut microbiome variability. PLoS Neglected Tropical Diseases 11, e0005677.

Slavic, K., Delves, M.J., Prudêncio, M., Talman, A.M., Straschil, U., Derbyshire, E.T., Xu, Z., Sinden, R.E., Mota, M.M., Morin, C., et al. (2011). Use of a selective inhibitor to define the chemotherapeutic potential of the plasmodial hexose transporter in different stages of the parasite’s life cycle. Antimicrobial Agents and Chemotherapy 55, 2824–2830.

Stone, W.J.R., Eldering, M., van Gemert, G.-J., Lanke, K.H.W., Grignard, L., van de Vegte-Bolmer, M.G., Siebelink-Stoter, R., Graumans, W., Roeffen, W.F.G., Drakeley, C.J., et al. (2013). The relevance and applicability of oocyst prevalence as a read-out for mosquito feeding assays. Scientific Reports 3, 272.

Takken, W., Smallegange, R.C., Vigneau, A.J., Johnston, V., Brown, M., Mordue-Luntz, A.J., and Billingsley, P.F. (2013). Larval nutrition differentially affects adult fitness and Plasmodium development in the malaria vectors Anopheles gambiae and Anopheles stephensi. Parasites & Vectors 6, 345.

Thadani, R., and Tammi, M.T. (2006). MicroTar: predicting microRNA targets from RNA duplexes. BMC Bioinformatics 7 Suppl 5, S20.

Wang, X., Hou, Y., Saha, T.T., Pei, G., Raikhel, A.S., and Zou, Z. (2017). Hormone and receptor interplay in the regulation of mosquito lipid metabolism. Proceedings of the National Academy of Sciences of the United States of America 201619326.

Yoon, M.-S. (2016). The Emerging Role of Branched-Chain Amino Acids in Insulin Resistance and Metabolism. Nutrients 8.

Zhang, S., Zeng, X., Ren, M., Mao, X., and Qiao, S. (2017). Novel metabolic and physiological functions of branched chain amino acids: a review. Journal of Animal Science and Biotechnology 8, 10.

Zhou, G., Flowers, M., Friedrich, K., Horton, J., Pennington, J., and Wells, M.A. (2004a). Metabolic fate of [14C]-labeled meal protein amino acids in Aedes aegypti mosquitoes. Journal of Insect Physiology 50, 337–349.

Zhou, G., Pennington, J.E., and Wells, M.A. (2004b). Utilization of pre-existing energy stores of female Aedes aegypti mosquitoes during the first gonotrophic cycle. Insect Biochemistry and Molecular Biology 34, 919–925.

Ziegler, R., and Ibrahim, M.M. (2001). Formation of lipid reserves in fat body and eggs of the yellow fever mosquito, Aedes aegypti. Journal of Insect Physiology 47, 623–627.

